# Uncertainty propagation in matrix population models: gaps, importance, and guidelines

**DOI:** 10.1101/2023.08.25.554793

**Authors:** Emily G. Simmonds, Owen R. Jones

**Affiliations:** Institutt for Biologi, Norwegian University of Science and Technology, Trondheim, Norway; Institute of Ecology and Evolution, School of Biological Sciences, University of Edinburgh, Edinburgh, UK; Department of Biology, University of Southern Denmark, Odense C, Denmark

**Keywords:** Matrix population models, uncertainty, propagation, simulation, population modelling

## Abstract

1. Matrix population models (MPMs), which describe the demographic behaviour of a population based on age or stage through discrete time, are popular in ecology, evolutionary biology, and conservation biology. MPMs provide a tool for guiding management decisions and can give insight into life history trade-offs, patterns of senescence, transient dynamics, and population trajectories. These models are parameterised with estimates of vital rates (e.g. survival and reproduction) and can have multiple layers of underlying statistical analyses, all of which introduce uncertainty. For accurate and transparent results, this uncertainty should be propagated through to quantities derived from the MPMs. However, full propagation is not always achieved, leading to omitted uncertainty, and negative consequences for the reliability of inferences drawn.
2. We summarised the state-of-the-art regarding vital rate uncertainty reporting and propagation, by reviewing papers using MPMs from 2010-2019. We then used reported uncertainties as the basis for a simulation study to explore the impact of uncertainty omission on inferences drawn from the analysis of MPMs. We simulated four scenarios of vital rate propagation and evaluated their impact on population growth rate estimates.
3. Although around 78% of MPM papers report some kind of uncertainty in their findings, only half of those report uncertainty in all aspects. Additionally, only 31% of papers fully propagate uncertainty through to derived quantities. Our simulations demonstrate that, even under a median uncertainty level, incomplete propagation introduces bias. Omitting uncertainty may substantially alter conclusions, particularly for results showing small changes in population size. Biased conclusions were most common when uncertainty in the most influential vital rates for population growth were omitted.
4. We suggest comprehensive guidelines for reporting and propagating uncertainty in MPMs. Standardising methods and reporting will increase the reliability of MPMs and enhance the comparability of different models. These guidelines will improve the accuracy, transparency, and reliability of population projections, increasing our confidence in results that can inform conservation efforts, ultimately contributing to biodiversity preservation.

## Introduction

Uncertainty is a fundamental part of the scientific process. Uncertainty is particularly relevant in statistical modelling and is an inherent part of most ecological models. This uncertainty can profoundly affect inferences drawn from these models, and therefore, the quantification and reporting of uncertainty is a fundamental and increasingly recognised part of robust science, particularly for predictive modelling (Dietze, Fox, Beck-Johnson, et al., 2018; Harwood & Stokes, 2003; Milner-Gulland & Shea, 2017). Predictive modelling can have many sources of uncertainty because there are typically many modelling steps taken and multiple models used to generate predictions. Each step and each model include some degree of uncertainty that can influence outcomes and should be propagated throughout the modelling process. When insights from such models are used to inform action, as exemplified by the use of predictive models during the COVID-19 pandemic (Edeling, Arabnejad, Sinclair, et al., 2021; James, Salomon, Buckee, et al., 2021; Kreps & Kriner, 2020; Meehan, Rojas, Adekunle, et al., 2020) or in fisheries management (Aeberhard, Flemming, & Nielsen, 2018; Harwood & Stokes, 2003; Kindsvater, Dulvy, Horswill, et al., 2018), the consequences of an incorrect or incomplete consideration of uncertainty include economically costly sub-optimal action (Aeberhard, Flemming, & Nielsen, 2018), damaged credibility (Kreps & Kriner, 2020; Palliser & Dodson, 2017), or even loss of life (Edeling, Arabnejad, Sinclair, et al., 2021; Regan, Ben-Haim, Langford, et al., 2005). Despite these potentially severe consequences, model-related uncertainties are not yet consistently or completely considered in ecological studies (Simmonds, Adjei, Andersen, et al., 2022).

Matrix population models (MPMs) (Caswell, 2001; Lefkovitch, 1965; Leslie, 1945) are widely used in predictive ecological research and have been created for at least 792 plant and 429 animal species (Salguero-Gómez, Jones, Archer, et al., 2016, 2015). MPMs capture how individuals, organised by age or stage, contribute to changes in population size and structure via the vital rates of survival, ontogenetic development, and reproduction through discrete time. The heart of the MPM is a square projection matrix (***A***) of estimated vital rates, which is used to project a population vector (**n**). MPMs are often composite models because the elements of the ***A*** matrix are estimated using statistical models or calculations (such as survival analysis, linear models, or calculations of central tendency). Each vital rate estimate has associated uncertainty because they are typically based on imperfect data from a sample of the population of interest (Caswell, 2001). Consequently, to achieve accurate uncertainty estimates for derived quantities, projections, and comparative analyses, uncertainty in the vital rates should be propagated through the modelling process. Despite the foundational recognition of the role of uncertainty quantification and propagation in drawing inferences from MPMs (see (Caswell, 2001, p 300), whether uncertainty is consistently and appropriately handled in practice remains an open question.

MPMs are not a homogenous group of models. The data used to underpin them can vary substantially, both within and between studies, and so do approaches to modelling vital rates. This methodological variation presents a potential source of inconsistent or incomplete quantification of uncertainty. While some studies collect the data needed to estimate every vital rate in the ***A*** matrix for the population under study (e.g. Bjørkvoll, Lee, Grøtan, et al., 2016), others consolidate their own estimates with estimates derived from other sources. These other sources are highly variable: estimates may be from the same population at a different time (Haslob, Hauss, Petereit, et al., 2012), from the same species at a different location/time (Stratford, Pollino, & Brown, 2016), from closely related species, or even “guestimates” from experts (Beissinger, 2014). Some data sources, particularly those relying on expert opinion or literature estimates, can challenge uncertainty quantification and cause omissions (e.g. Gamelon, Besnard, Gaillard, et al., 2011). A further barrier to complete uncertainty reporting is that different models are often used to estimate reproduction and survival processes (e.g. Ali, Kauffman, Amin, et al., 2018; Carson, Levin, Cook, et al., 2011; Paez, Bock, Espinal-Garcia, et al., 2015). Such differential methodological practices and different data availability could lead to missing uncertainty estimates for particular stages, ages, or vital rates; for example, uncertainty may be reported for a mark-recapture survival analysis but not for mean reproductive success (Paez, Bock, Espinal-Garcia, et al., 2015). Omissions of key model-related uncertainties are common in ecology and evolution (Simmonds, Adjei, Andersen, et al., 2022) and we might expect the same within MPM studies.

It is also important for models to fully account for how uncertainty in lower-level parameters propagates through the model’s calculations to influence higher-level estimates. However, the full propagation of vital rate uncertainty into derived estimates and projections from MPMs is prevented if uncertainty estimates are missing for any of the lower-level vital rates. The consequences of incomplete or absent uncertainty propagation are not restricted to the focal study. Meta-analyses and comparative studies that use estimates from MPMs will inherit any imperfections in constituent studies with potentially unanticipated consequences. Open access databases, such as COMPADRE (Salguero-Gómez, Jones, Archer, et al., 2015) and COMADRE (Salguero-Gómez, Jones, Archer, et al., 2016), have greatly contributed to the field of predictive ecological modelling by providing access to thousands of published MPMs and metadata for comparative analyses (Cant, Capdevila, Beger, et al., 2023; Enríquez, Daalen, & Caswell, 2022; Hernández-Yáñez, Kim, & Che-Castaldo, 2022; Lanuza, Rader, Stavert, et al., 2023). However, these repositories do not provide any uncertainty metrics for the underlying parameters of the matrices they hold or their derived quantities, causing uncertainty omission. Though uncertainty omission of this nature could lead to overconfident or biased inferences (Edeling, Arabnejad, Sinclair, et al., 2021), the magnitudes of such impacts are not yet known.

We currently lack a standardised protocol for uncertainty quantification, propagation, and reporting for MPMs. Recent efforts to develop a standard protocol for reporting MPMs (Gascoigne, Rolph, Sankey, et al., 2023) are vitally important and stress the importance of accompanying vital rate estimates with a metric of uncertainty. Although following this protocol will greatly aid interpretation and transparency, the protocol is insufficient for consistent propagation, comparison, or easy storage of uncertainty data in online databases. The inconsistent use of metrics in different studies (e.g. standard error vs confidence intervals), particularly when uncertainty distributions are not Normal, presents a challenge for comparing uncertainty across papers or propagating uncertainties into follow-on analyses. Consequently, additional uncertainty-focused guidelines are required to build on Gascoigne’s protocol and further improve the reporting of MPMs.

In this paper, we suggest best-practice guidelines for uncertainty reporting and propagation for MPMs, building on Gascoigne et al.’s (2023) protocol. To inform the development of these guidelines, we review MPM papers to identify any gaps/inconsistencies in the current consideration of uncertainty in MPMs. To highlight the need for such guidelines, we conduct a simulation study to show what happens when uncertainty is not propagated correctly. We ask two specific questions about the importance of uncertainty propagation for the robustness and reliability of conclusions drawn from MPMs: 1. How does propagation of uncertainty alter conclusions around population growth trajectories? 2. How does life history (semelparous vs iteroparous and reproductive dominant vs survival dominant) influence the impact of partial propagation of vital rate uncertainty?

## Method

### Review

To review how often vital rate uncertainties were reported and propagated in papers using MPMs, we adopted a systematic, but not exhaustive, approach to paper selection. We focussed specifically on animal studies and used the open source COMADRE database (“COMADRE Animal Matrix Database,” 2021, v.4.21.8.0, accessed February 2023). We extracted matrices from articles with a DOI that had a publication date from 2010-2019. We focused on the most recent decade to ensure we reviewed the state-of-the-art rather than historical trends. The use of DOI restricted the sample to studies that had gone through peer review, though this was not a perfect metric (see below). We excluded studies with less than one year duration and matrices with missing values, leaving 121 publications to review. Three of these publications were removed due to lack of access, as they were behind a inaccessible paywall. A further two were removed as they were not peer reviewed; one thesis and one technical government report. We classed theses as non-peer reviewed as it is impossible to be sure if a thesis is published before or after review by examiners. A further 11 papers were not reviewed as they did not contain an MPM. This final filter left 105 papers for review (see Supporting Information, Section S10).

To conduct the review and assess uncertainty reporting, quantification, and propagation, we answered 13 questions covering model structure, uncertainty details, species, and sample size for each paper (Table 1). Where papers included different treatments of uncertainty across several MPMs, we took a conservative approach and answered based on the most comprehensively reported matrix. In addition, we recorded vital rates and any associated uncertainty and noted whether the uncertainty was propagated and how. For uncertainty type, it should be noted that standard deviation indicates the standard deviation of a sampling distribution or posterior sample, not a standard deviation of observed data.

**Table 1:** Questions and descriptions used as the basis of the review of the state-of-the-art for uncertainty reporting in MPMs between 2010-2019. Italics indicate answer descriptions.

| # | Question | Answer1 = primary answer to the question | Answer2 = further details that explain or support first answer | DOI = the DOI of the paper | Year = Year the paper was published | Author = first author last name (as defined by journal) | Comments |
| --- | --- | --- | --- | --- | --- | --- | --- |
| 1 | Was there an MPM in original paper? | *yes = fill in rest of table*<br>*no = exclude* | *write what model is used if Answer 1 = no* |  |  |  |  |
| 2 | What was the focal organism? | *The order the study species belongs to* | *species (common or Latin name)* |  |  |  |  |
| 3 | Did they model an explicit observation process for survival? | *Yes = something like capture-recapture model or direct calculation i.e. 5/10*<br>*no = just logistic regression* | *Put details of the model or calculation used here*<br>*or "from literature"* |  |  |  |  |
| 4 | Did they model an explicit observation process for reproduction? | Yes = include extra count error<br>no = standard lm or glm or direct calculation | Put details of the model or calculation used here<br>or "from literature" |  |  |  |  |
| 5 | Was uncertainty reported? | Yes, no, maybe, NA, can't tell (this includes any uncertainty in derived quantities e.g. lambda) | Add details of which elements uncertainty IS reported for |  |  |  |  |
| 6 | If reported, was the uncertainty complete for all parameters? | Yes, no, NA, can't tell (here focus is only on matrix elements/transitions NOT derived quantities) | If no – list those that it is missing for (just broad level e.g. adult survival, reproduction etc) |  |  |  |  |
| 7 | Was lambda estimated from the matrix? If yes what was the conclusion | yes, no, NA, can't tell | If yes – write conclusion here |  |  |  |  |
| 8 | Was a sensitivity/elasticity analysis/LTRE conducted? If yes what was the conclusion? | yes, no, NA, can't tell | If yes – write conclusion here |  |  |  |  |
| 9 | What was the census type? | pre, post, period/continuous birth | More detail if necessary i.e. where the information is from |  |  |  |  |
| 10 | Was the life cycle correctly estimated? | yes, no, can't tell<br>For details of potential errors see:<br>https://www.sciencedirect.com/science/article/pii/S0304380019301085 | More detail if necessary i.e. where the information is from |  |  |  |  |
| 11 | What was the error metric (if found)? | Write error here or NA | More detail if necessary i.e. where the information is from |  |  |  |  |
| 12 | What metric was used for reproduction? | Do they use recruits, fledgling etc or do they use a compound metric e.g. clutch size • nest survival • juvenile survival | More detail if necessary i.e. where the information is from |  |  |  |  |
| 13 | What was the sample size? | maximum reported sample size | What did it represent i.e. total caught, total in reproduction etc |  |  |  |  |

**Table 2:** Table of our recommended guidelines for uncertainty reporting and propagation in MPMs alongside the rationale for each guideline.

| Number | Guideline | Rationale |
| --- | --- | --- |
| 1 | **Report uncertainty for all matrix elements (following general protocol in (Gascoigne, Rolph, Sankey, et al., 2023))** | Complete reporting of uncertainties gives the full, transparent picture of the uncertainty associated with the MPM. It also gives necessary resources for full propagation. |
| 2 | **Report uncertainty as a standard error (or standard deviation of a posterior or sampling distribution) coupled with a 95% interval (confidence, credible, or quantile) on the same scale as the estimate** | Both a measurement of error and an interval are required for ease of interpretation and reuse. Propagation of uncertainty through parametric resampling requires a standard error or equivalent. In contrast, 95% intervals can be more immediately interpretable and aid cross-study comparisons of results. |
| 3 | **Include details of the assumed distribution of the target parameter or its associated uncertainty distribution** | Necessary to propagate uncertainty to further modelling steps, for example through parametric resampling |
| 4 | **Report sample sizes for each vital rate, not just the study as a whole** | Required to generate some estimates of sampling uncertainty |
| 5 | **Report uncertainty numerically as a minimum. This can be complemented with visual presentation to aid interpretation** | Numeric presentation is required for accurate reuse of results in either meta-analyses or follow on studies and for unambiguous interpretation |
| 6 | **Include all data and code as minimum and resampling results, where possible, with the published paper. This can be achieved either as supporting** | Essential for full replication or reuse of results. In particular, Bayesian MCMC sampling or various forms of bootstrap resampling are time consuming and |
|  | information or ideally archived in its own preserved repository with a DOI | computationally demanding analysis steps but are required to obtain a full estimate of uncertainty distributions. For many demographic parameters their uncertainty distribution is not normally distributed, as shown in Figures 2 and 3. As a result, summary metrics such as a standard error or credible interval will not fully capture the uncertainty distribution. Therefore, providing the full set of samples will aid reuse and further propagation of the uncertainty. |

To analyse the results from our review, we calculated the percentage of papers that fell into each of the following categories: reported any uncertainty (i.e. answered ‘yes’ to question 5), reported uncertainty for all vital rates (i.e. answered ‘yes’ to question 6), missed all vital rate uncertainty (i.e. answered ‘no’ to question 5 and had both ‘survival’ and ‘reproduction’ noted in ‘Answer2’ question 5), missed uncertainty for survival (i.e. answered ‘no’ to question 5 and had ‘survival’ noted in ‘Answer2’ question 5), missed uncertainty for reproduction (i.e. answered ‘no’ to question 5 and had ‘reproduction’ noted in ‘Answer2’ question 5), reported uncertainty and propagated it (i.e. answered ‘yes’ to question 5 and then the percentage of vital rate uncertainties that were propagated).

### Simulation of uncertainty propagation

We used simulations to assess the importance of propagating vital rate uncertainty to derived estimates. As the basis of our simulations, we used the same sample of 2,123 matrices from 121 studies matrices that we used in our review. Although this sample included a mixture of stage-based and age-based matrices, we opted to treat them all similarly for this analysis. Our rationale for this is that, because our analysis only relies on the linear transformation of the matrices, treating age-based matrices as stage-based would not alter our results: i.e. the dominant eigenvalue, *λ*, would be the same whether a matrix is age- or stage-based. For simplicity, we hereafter refer to all MPMs in our sample as stage-based. We divided this sample of matrices into six groups based on matrix dimension (three groups) and reproductive strategy (the distribution of reproduction across stages, two groups). The groups for matrix dimension, were two-by-two (n = 168), three-by-three (n = 218), and five-by-five (n = 92) matrices. For reproductive strategy, we defined two groups: those that reproduce in only a single stage and those that reproduce in several stages. For the two-by-two and three-by-three matrices, we defined “several stages” as reproducing in all stages, but for five-by-five matrices, we defined it as reproducing in three or more stages. Hereafter we refer to these six groups as dimension-by-strategy groups.

To ensure that our simulations covered a large range of life histories, we stratified these six dimension-by-strategy groups by sorting the matrices within each group based on the ratio of elasticities for reproduction and survival. We deemed MPMs with a ratio of <1 to be *survival dominant* and those with a ratio of >1 to be *reproduction dominant*. We then selected five MPMs from each of the six dimension-by-strategy groups at approximately the 12.5th, 25th, 50th, 75th and 87.5th percentiles of this ratio (see Supporting Information, Section S9). We made one exception to this methodology for the five-by-five and reproduce once group. For this group, the matrix selected for the 12.5th percentile had a survival value of 1 for one stage, which is impossible in reality. Therefore, we selected the preceding MPM sequentially (based on reproduction:survival ratio). By following this stratified sampling approach, we obtained 30 MPMs that we could use as base MPMs for our simulations.

Next, to explore how uncertainty affects inferences from MPMs, we added uncertainty to every vital rate in each base MPM. Rather than use an arbitrary degree of uncertainty, we obtained “realistic” uncertainty measures for two broad types of vital rate (reproduction-related and survival-related) from the literature. We did this by examining the reported uncertainties from our uncertainty review and sorted them into ‘reproduction’ (contribution to the youngest stage) or ‘survival’ (remaining in stage or transitioning to another stage) elements using matrix position or vital rate name from the original paper wherever possible. To facilitate comparability of uncertainty across studies and mitigate the influence of the specific uncertainty metric on our analyses, we exclusively employed confidence intervals and standard errors as measures of uncertainty. We converted confidence intervals back into standard errors for their application in our simulations (see Supporting Information, Section S1).

We classified the type of vital rate as either reproduction-or survival-related by considering their position in the matrix and their names in the original publication. Regarding matrix position, we deemed elements in the first row to be reproduction-related. In addition, we assumed that rates referred to as “reproduction”, “recruitment”, “fecundity”, and “fertility” were all reproduction-associated, while those termed “survival” and “stasis” were survival-related. Using these heuristics, it is possible that some retrogression could be included as reproduction or survival, but we expect this to be minimal. If we could not determine whether a vital rate’s uncertainty was reproduction- or survival-related, we omitted it from our analysis. After collecting these reported uncertainty measures, we defined low-, mid- and high-level uncertainty as the 2.5^th^, 50^th^ and 97.5th percentile for each vital rate type.

We used these 30 base MPMs, for the three levels of uncertainty (i.e. 90 combinations), to investigate how uncertainty affects results by conducting parametric resampling of each matrix element. For the resampling, we drew values from a log-normal distribution for the reproductive parameter (*λ*_*r*_) using the `lnorms()` function from the `popbio` package (Stubben & Milligan, 2007), which is the rate parameter from a Poisson distribution. For survival, we drew values from a beta distribution to resample the probability parameter from a Binomial distribution using the `betaval()` function from the `popbio` package. For each resampling, we determined the mean from the base matrix vital rate and the variance or standard deviation of the distribution using one of the uncertainty levels extracted from the reviewed papers. We included uncertainty as a proportion of the base estimate, rather than an absolute value, to account for the different absolute magnitude of the vital rates in each reviewed paper and in the base matrices for the simulations. We performed 10,000 resamples across all parameters simultaneously. We then propagated uncertainty from the vital rates to estimates of *λ* using this resampling under four scenarios:

1. No uncertainty - this will be the output from the original base matrix.
2. Reproduction only - this is where reproductive elements were resampled, but survival elements were not.
3. Survival only - survival elements were resampled, but reproductive elements were not.
4. Full propagation - all elements were resampled at once.

To assess how these scenarios impacted MPM results, we estimated two derived metrics for each resampled matrix: estimated population growth rate (*λ*) (the dominant eigen value of the matrix, calculated using the `eigs()`function from the `popdemo` package (Stott, Hodgson, Townley, et al., 2021)) and the dominant elasticity (relative sensitivity), giving 10,000 estimates of each performance metric per base matrix. To determine if conclusions were changed, we characterised our results into three categories: stable (*λ* > 0.98 < 1.02), increasing (*λ* > 1.02), decreasing (*λ* < 0.98). We then calculated how many of the resampled *λ* values matched the original conclusion obtained from the base matrix.

## Results

Across all scenarios, matrix dimension had a minimal influence on results, causing no discernible difference in uncertainty distributions under the different propagation scenarios. Conclusion changes showed some differences for the five-by-five matrices, where conclusions were more likely to change under uncertainty propagation for species that reproduce in multiple stages, but less likely for species that reproduce in only one stage. Below we present results for the three-by-three matrices under the mid-level uncertainty, all other results can be found in the Supporting Information, Sections S1-2, S5-8.

### Uncertainty reporting is common, but gaps remain

From our review of uncertainty reporting and propagation in the MPM literature, we found that although a healthy 78% of papers report some uncertainty, this reporting is incomplete (i.e. some vital rates do not include uncertainty estimates) for 50% of those papers. Of the 41 papers with incomplete reporting, nine (22%) were missing reproduction uncertainty, eight (20%) were missing survival uncertainty, and 23 papers (56%) were missing both reproduction and survival uncertainty, only reporting uncertainty in derived quantities (the remaining paper had unclear reporting). 56% of papers that reported uncertainty propagated at least some of it to derived estimates (Figure 1). Incomplete reporting coupled with non-perfect propagation resulted in just under a third of papers that achieved complete propagation of vital rate uncertainties. As is shown in Figure 1, partial propagation can arise from three processes; complete reporting of uncertainties in vital rates but incomplete propagation (<1% of papers), incomplete reporting of uncertainties in vital rates and incomplete propagation (3% of papers), and incomplete reporting of uncertainties in vital rates and complete propagation (9% of papers). It should be noted that we still class this latter process as partial propagation, even though it represents the propagation of all quantified/reported uncertainty. In total, it was impossible to ascertain whether propagation occurred in 22% of papers.

**Figure 1:**
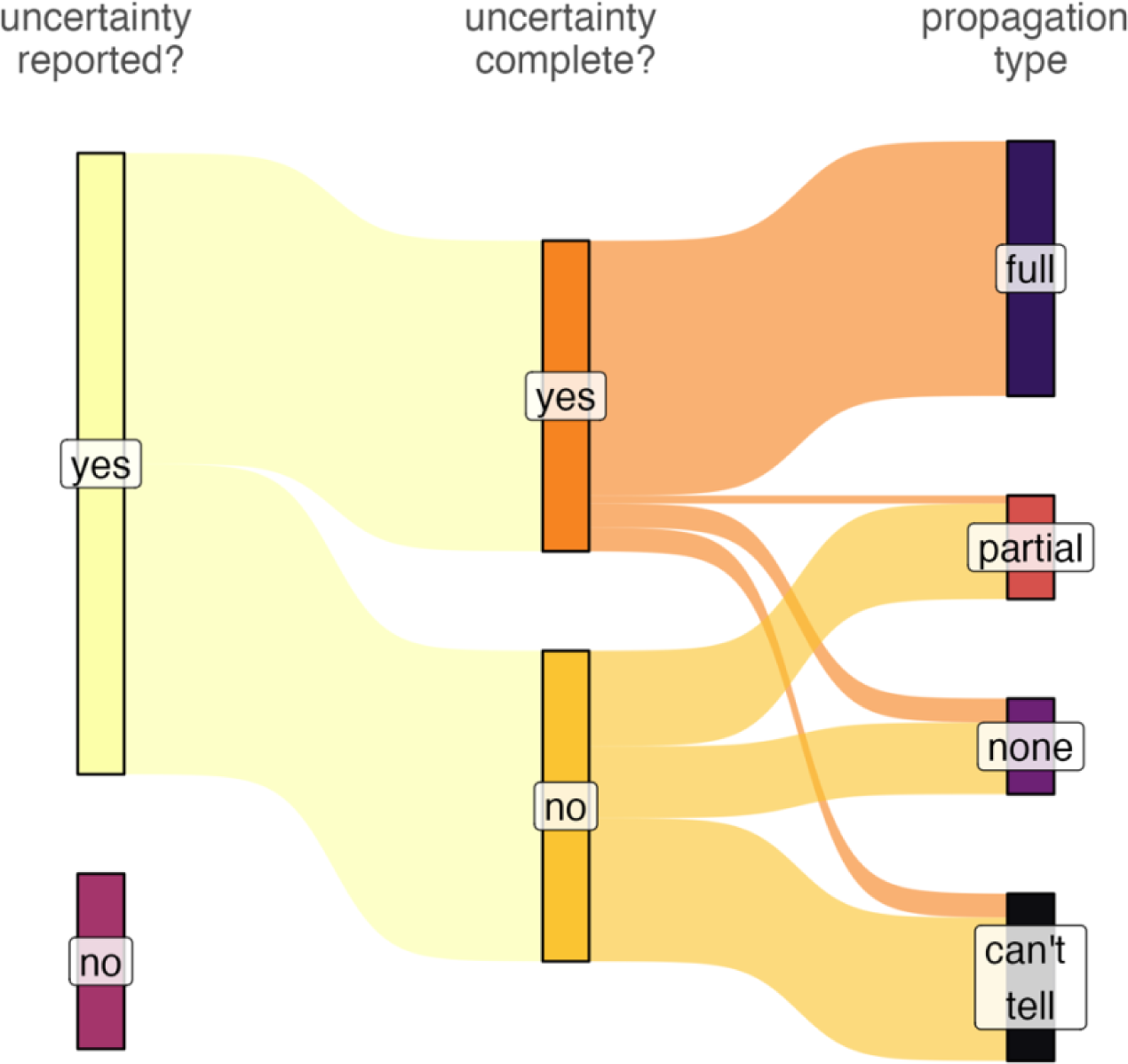
Results of our review of uncertainty reporting and propagation in recent matrix population modelling papers. The thickness of the flow segments corresponds to the proportion of reviewed papers that fall in the highlighted category. These results show how uncertainty is omitted at various points during the modelling process leading to a large proportion of papers failing to achieve full uncertainty propagation.

### Life history strategy influences the consequences of partial propagation

The results of our parametric resampling analysis indicate a consistent pattern across propagation scenarios and reproductive strategies (Figure 2). Specifically, as the dominance of reproduction (as quantified by the ratio of elasticities for reproduction and survival) in the MPM increases, so does the importance of propagating the uncertainty in reproduction for achieving the full uncertainty distribution of *λ*, indicated by the full propagation scenario. For example, for panels e and j in Figure 2, indicating results for the MPMs with the highest reproduction:survival ratio, distribution of λ in the ‘reproduction only’ scenario reflects the distribution from the ‘full propagation’ scenario very well. Despite this similarity of the λ distributions for partial and full propagation in some cases, the overall pattern is that partial propagation tends to underestimate the uncertainty in λ. For most scenarios presented in Figure 2, large variability leads to results spanning a *λ* of 1, indicating no clear population trajectory. There are some exceptions to this pattern, such as panels b and j where the populations are estimated to increase in almost all resamples.

**Figure 2:**
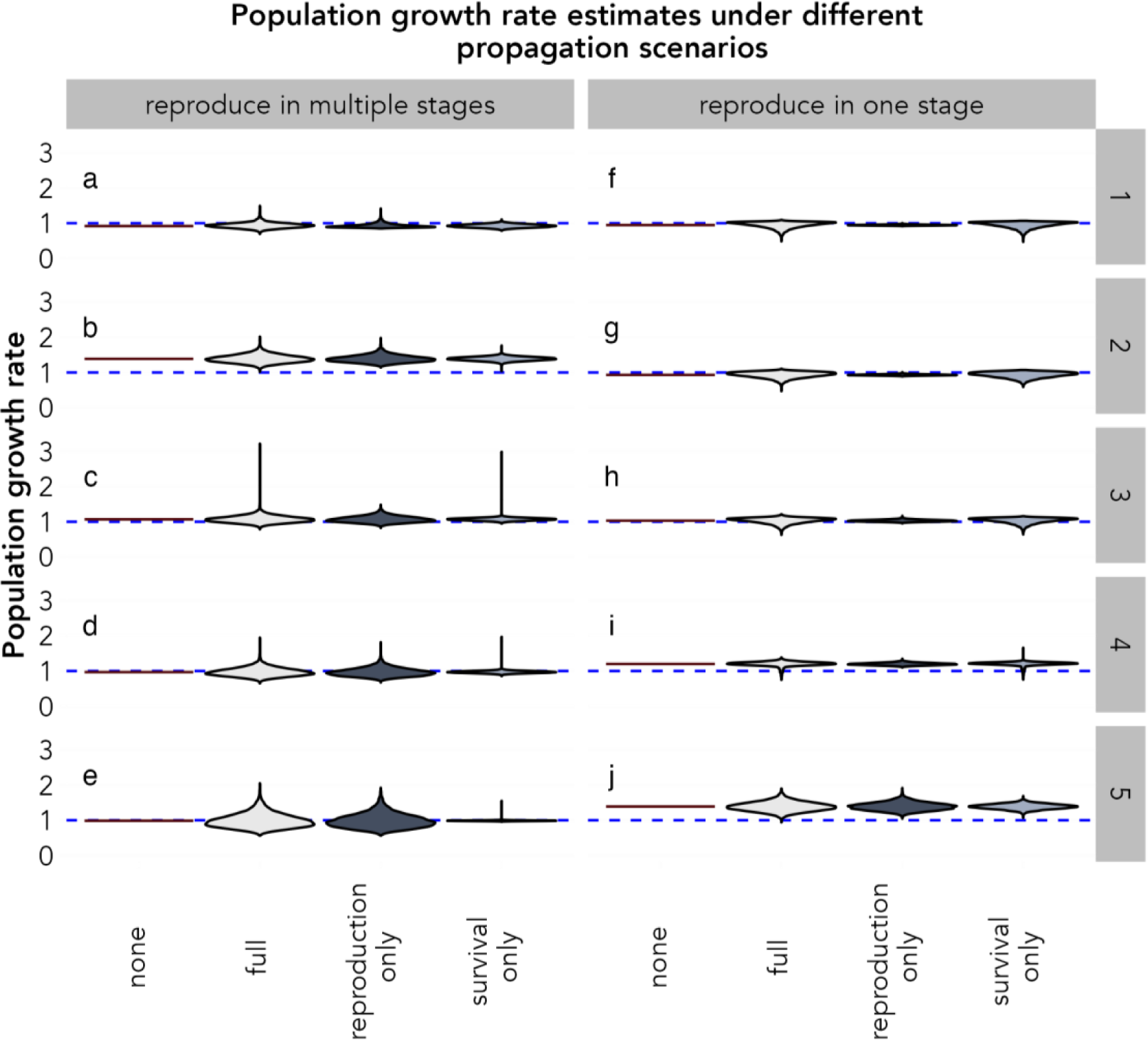
Population growth rate distributions under different types of uncertainty propagation and with different life histories, estimated using parametric resampling. The left-hand column (panels a-e) shows cases where reproduction occurs in multiple stages, while the right-hand column (panels f-j) shows cases where reproduction occurs in a single stage only. The relative importance of reproduction vs survival changes systematically from rows 1 to 5 such that row 1 is relatively survival dominant and row 5 is relatively reproduction dominant. The blue dashed line indicates a population growth rate of 1. All results are for a three-by-three matrix at the mid-uncertainty level. This figure shows how bias caused by partial propagation shifts with the life history of an organism.

### Failure to propagate uncertainty can lead to incorrect conclusions

Parametric resampling changed the conclusion one would draw about population trajectory (increasing, decreasing, or stable) (Figure 3). Most of our selected MPMs showed substantial changes in their conclusion depending on whether uncertainty was fully or partially propagated. Notable exceptions are panels b, i, and j, for which the conclusion showed no or minimal changes. These instances of minimal change had estimated *λ* values from the base matrix that were furthest from 1, suggesting strongly increasing or decreasing trajectories are not substantially changed by mid-level uncertainty propagation. The bias induced by partial propagation can be seen in Figure 3 panels f and g, where propagating uncertainty only in reproduction led to the same conclusion as the base matrix. However, conclusion changes arose when uncertainty in survival was propagated, either alone or as part of full propagation. Overall, propagating uncertainty changed conclusions about population trajectory in seven of the 10 life history strategies tested. This rate of 70% is higher than the rate found in our review, where 42.5% of *λ* estimates with reported uncertainty had confidence spanning 1. However, as our review did not assess the impact of uncertainty for papers that did not report it, 42.5% likely underestimates the true percentage.

**Figure 3:**
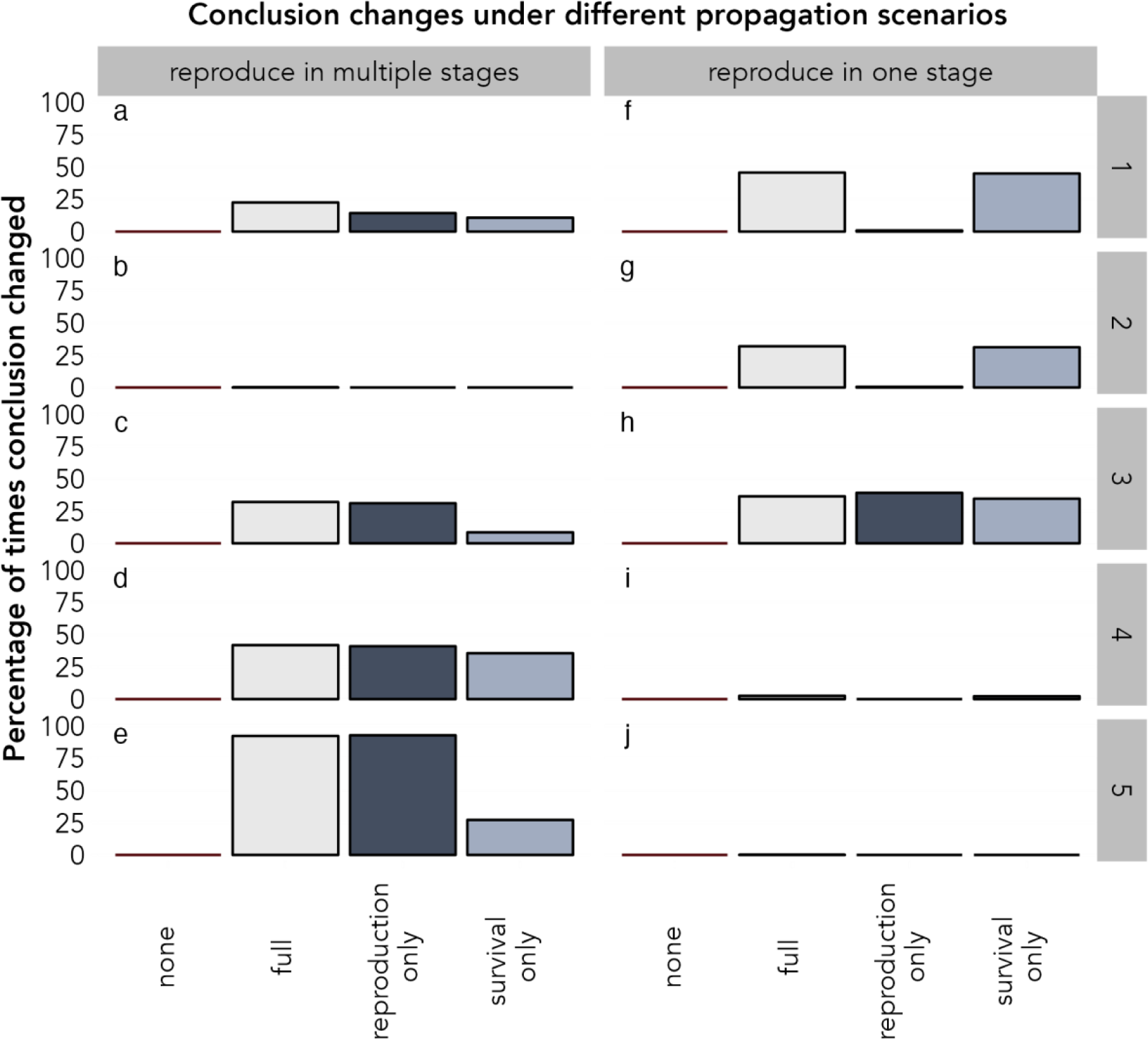
The percentage of times a conclusion changed during parametric resampling under different types of uncertainty propagation and with different life histories. The left-hand column (panels a-e) shows cases where reproduction occurs in multiple stages, while the right-hand column (panel f-j) shows cases where reproduction occurs in a single stage only. The relative importance of reproduction vs survival changes systematically from rows 1 to 5 such that row 1 is relatively survival-dominant, and row 5 is relatively reproduction-dominant. All results are for a three-by-three matrix at the mid-uncertainty level. This figure shows how a failure to fully propagate uncertainty can alter conclusions of population trends.

**Figure 4:**
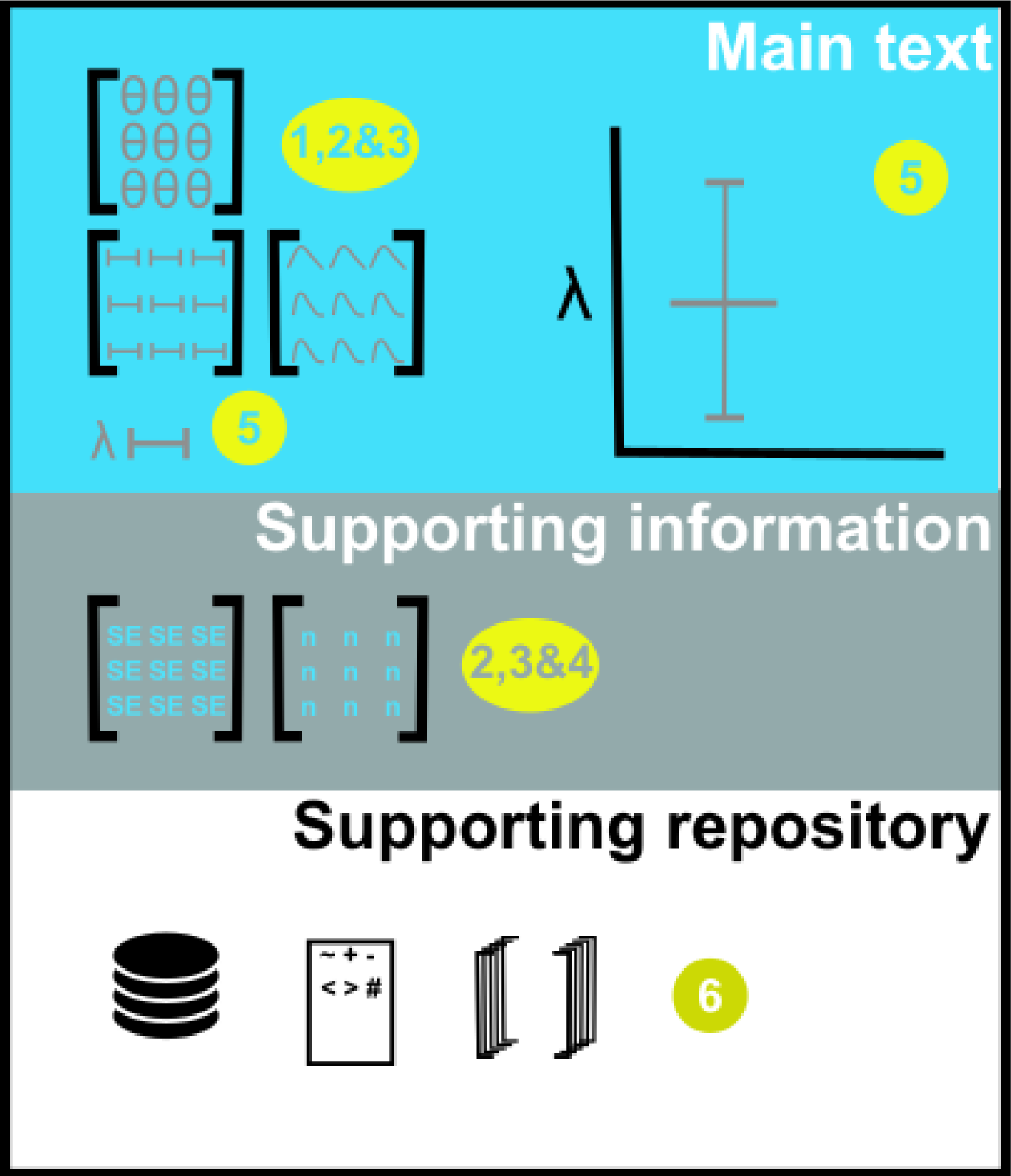
Illustration of implementation of our guidelines. The numbers in yellow circles map onto our numbered guidelines. Each section indicates a different location within a manuscript. The main text section illustrates the presentation of mean estimates for each vital rate in a MPM, the associated confidence/credible interval, the assumed distribution of the uncertainty, and numerical and visual presentation of a derived quantity (*λ*). The supporting information section illustrates the presentation of standard errors (or standard deviation of a posterior distribution) and sample size for each element of an MPM. The supporting repository section illustrates presentation of the data, code, and resamples (Bayesian, MCMC, or bootstrap).

In contrast to the results for *λ*, the identity of the dominant vital rate was largely robust under uncertainty propagation at low uncertainty levels and for smaller matrices (two-by-two): the vital rate identified as dominant changed in fewer than 5% of the full propagation resamples (see Supporting Information, Section S4). In contrast, larger matrices, such as the five-by-five, and higher uncertainty levels (mid and high) produced greater variability. In particular, the five-by-five matrices for semelparous species showed large changes in elasticity with up to 100% of resamples estimating a different dominant vital rate to the base matrix estimate (see Supporting Information, Section S4).

## Discussion

Uncertainty is a key element of the scientific process but one that can often be overlooked (Simmonds, Adjei, Andersen, et al., 2022), with potentially detrimental consequences (Aeberhard, Flemming, & Nielsen, 2018; Edeling, Arabnejad, Sinclair, et al., 2021; Regan, Ben-Haim, Langford, et al., 2005). In this study we explored the state-of-the-art of uncertainty reporting and propagation in papers using MPMs, as well as the consequences of omission. Our results show that while the reporting of vital rate uncertainty is common for MPMs, it is inconsistent across the subfield and often incomplete. We show that neglecting uncertainty, particularly during error propagation, can lead to erroneous conclusions regarding population trajectories. The degree of impact from missing or partial uncertainty propagation depends on the life history of the organism being studied and its population trajectory.

### Areas for improvement in uncertainty reporting

The high prevalence of reporting at least some uncertainty in MPM papers (78%) is a commendable achievement. This subfield performs substantially better at reporting parameter-related uncertainty than other areas in ecology (Simmonds, Adjei, Andersen, et al., 2022). Nevertheless, this reporting prevalence is still lower than in other fields such as political or health science (Simmonds, Adjei, Andersen, et al., 2022) and, therefore, there is room for improvement, as a fifth of MPM papers report no vital rate uncertainty at all.

One key area for improvement is in the completeness of uncertainty reporting across all vital rates. Currently, over half of papers omit at least one demographic process from their uncertainty consideration. This leads to an incomplete picture of uncertainty and can cause biased/overconfident results (Figures 2 and 3). Achieving complete uncertainty reporting is, however, not straightforward. Often, estimates for particular vital rates are obtained from the literature or are based on expert opinion due to a lack of data from which to estimate them statistically. This data deficiency is particularly common for species with cryptic life-cycle stages and for rare species (Beissinger, 2014; Fitzgerald, Smith, Culver, et al., 2021; Martin, Delheimer, Moriarty, et al., 2022). Yet the use of literature or expert estimates does not preclude the inclusion of uncertainty entirely. Although obtaining a more formal statistical estimate may be prohibitively challenging, other methods, such as defining a plausible range for a given vital rate, are available and have been implemented successfully in previous studies (Beissinger, 2014). These alternative methods are not without fault: They will be more subjective than estimating uncertainty statistically and have little foundation to assess their accuracy in capturing uncertainty. Nevertheless, we argue that including a ‘best guess’ of uncertainty is preferable to ignoring it completely.

Another key area for improvement is the consistency of reporting. There are numerous ways in which uncertainty is reported for MPMs, from confidence intervals, standard errors, and credible intervals, to process variance, standard deviations, and more. It is not always clear from reporting whether the uncertainty is presented on the scale of the estimate it is associated with or which distribution it is representing. Furthermore, several papers presented uncertainties only visually (Ali, Kauffman, Amin, et al., 2018; Hanly & Haase, 2016; Rota, Millspaugh, Rumble, et al., 2014), making it almost impossible to propagate this uncertainty to further studies. This inconsistent reporting mirrors recent findings about MPM reporting more broadly (Gascoigne, Rolph, Sankey, et al., 2023) and makes it challenging to compare uncertainty across studies, therefore limiting the potential to use it in comparative analyses. Inconsistency also brings challenges for incorporating uncertainty into open-access databases, as if it is not reported in a standardised way, it is not straightforward to store it in a standardised and interpretable way.

### Factors affecting uncertainty impacts

The consequences of uncertainty for downstream inference depend on the relative importance of reproduction vs survival and the population’s trajectory. To provide a full and transparent understanding of a study’s results, it is necessary to propagate all vital rate uncertainties into derived estimates. However, the amount of bias introduced by partial propagation varies depending on life history, particularly on the balance between the influence of reproduction and survival on population growth rate (*λ*) (Figure 2). Partial propagation tends to produce less biased and overconfident estimates when omitting uncertainty in the non-dominant process (as identified using the reproduction:survival ratio), compared to omitting uncertainty in the dominant process. The correlation between this ratio and generation time (Supporting Information, Section S9), suggests that propagating uncertainty in reproductive rates is typically more important for species with fast life histories, while propagating uncertainty in survival rates is more important for species with slower life histories. Focusing uncertainty analysis on the vital rates that most influence population growth (the “dominant” process) is critical, as these dominant rates naturally have the greatest effect on estimates of *λ*. Knowledge of which process dominates could help identify existing studies likely to have biased or overconfident results due to incomplete uncertainty propagation.

In most cases we explored, full propagation was required to obtain a complete uncertainty distribution for population growth rate. Partial propagation can lead to incorrect conclusions (Figure 2a, c, e, f, g). These incorrect conclusions could have particularly stark consequences where results inform species conservation or management actions (Aeberhard, Flemming, & Nielsen, 2018; Regan, Ben-Haim, Langford, et al., 2005) and where acting incorrectly could be costly in terms of money or lives (Aeberhard, Flemming, & Nielsen, 2018; Edeling, Arabnejad, Sinclair, et al., 2021). It is worth noting that all vital rates in our simulations were subject to the same proportional amount of uncertainty. This limitation could have an impact on the results relating to partial propagation, for example, if reproduction is the dominant process in a matrix but it has much lower proportional uncertainty than survival, survival uncertainty might still be more influential for *λ* than reproduction. Moreover, the identification of the dominant demographic processes for a population could also be influenced by uncertainty. We found that the estimation of the dominant vital rate through elasticity analysis seems reasonably robust to uncertainty propagation under low and medium uncertainty levels for matrices smaller than five-by-five (see Supporting Information, Section S4), which is reassuring for population management applications. However, we still urge caution because under high uncertainty, particular reproductive strategies, and larger matrices (five-by-five) the dominant processes are shown to be highly variable (see Supporting Information, Section S4) which could lead to spurious inferences. Consequently, relying on partial propagation should only be done with extreme caution.

Population trajectory had the most substantial impact on whether the propagation of vital rate uncertainty changed the conclusions from those obtained from the mean matrix. For 70% of the three-by-three matrices we examined, accounting for uncertainty led to substantial changes in conclusions (Figure 3). However, for 30% of the matrices, little or no change occurred. These matrices with little change were those with more sharply increasing or decreasing populations. This suggests that for populations that are more rapidly changing, propagating uncertainty from vital rates is unlikely to alter their conclusions. This finding is reassuring for work on vulnerable populations that are in decline (IUCN, 2022) as it implies that their results are likely to be robust even without accounting for uncertainty. This does not mean that uncertainty should be disregarded in such populations. If we take a declining population as an example, identifying a decline is just one step in conservation efforts. Understanding the rate of the decline, the key demographic processes driving it, and how to mitigate it are all crucial for conservation action and all more informative with well quantified uncertainty bounds. Indeed, a decline of 20% is vastly different to a decline of 5%, even though both are population declines.

Matrix dimension had a relatively small influence on the consequences of uncertainty omission compared to life history attributes and population trajectory (see Supporting Information, Sections S2-3, S5-8). This suggests that a modeller’s decisions on the number of stages or ages to use has less influence on the propagation of vital rate uncertainty than the intrinsic biology of the species. However, because all vital rates of the same type had the same relative amount of uncertainty in our study, this may have reduced the impact of matrix dimension. In empirical studies, splitting the population into a larger number of groups will likely decrease the sample size for each group, consequently increasing the uncertainty.

This could have knock-on impacts for larger matrices, potentially making their results more uncertain. This should be explored explicitly. One area where matrix dimension did have an impact was in the estimates of a dominant vital rate through elasticity analysis. The larger the matrix, the more likely the dominant vital rate was to be variable under uncertainty propagation (see Supporting Information, Section S4). This is likely driven by the larger number of possible vital rates in a larger matrix but highlights the importance of considering propagation into all derived quantities, and not just population growth rates. This was rarely achieved in the studies we reviewed, but we did identify some published examples of good practice (Bjørkvoll, Lee, Grøtan, et al., 2016; Foster & Vincent, 2012; Meyer, Robertson, Chilvers, et al., 2015).

Overall, our results show the omission of uncertainty has important consequences for the accuracy and completeness of results which could affect other studies that rely on MPM results, e.g. comparative studies. These impacts are likely to be more important when uncertainty levels are higher (see Supporting Information, Sections S7-8), though more research is needed to explore these impacts specifically. Therefore, it should be an urgent priority to include measures of uncertainty in the vital rates that underpin MPMs in any collections of MPMs, such as COMPADRE (Salguero-Gómez, Jones, Archer, et al., 2015) and COMADRE (Salguero-Gómez, Jones, Archer, et al., 2016) and give them the same status as the estimates themselves (Caswell, 2001). To facilitate this effort, we distil our findings into specific guidelines on how to deal with and report uncertainty in MPMs.

### Guidelines for uncertainty reporting in MPMs

These six guidelines are designed to aid the MPM community in the consistent and complete reporting and propagation of vital rate uncertainty and should be considered a minimum standard.

### Limitations

As with all scientific work, our study is based on a series of assumptions which limit the interpretation of its output. We detail the major limitations and their likely impacts below.

- In our simulations, we used the same relative uncertainty across all vital rates of the same type, which is likely unrealistic. The actual levels of bias from partial propagation in any given study will vary depending on the exact balance of uncertainty across all vital rates.
- We did not explore the impact of uncertainty propagation on all derived quantities that can be calculated from MPMs. For logistical reasons, we focused on the most commonly reported derived quantities of population growth rate (*λ*) and elasticities. The impact on other quantities could differ from the results presented here and we recommend further specific exploration.
- While we did explore a range of life histories and across a range of matrix dimensions, neither of these axes were explored exhaustively.
- We did not assess whether the uncertainty calculation or propagation methods were correct or most appropriate to a given situation. Considering this would have involved some potentially subjective assessments and would take a more time per paper reviewed. However, it could alter some results and give new insight if there are common methodological failings in MPM papers (Kendall, Fujiwara, Diaz-Lopez, et al., 2019).
- We do not account for covariance in uncertainty among vital rates. For example, the uncertainty for each stage-based survival estimate may not be independent due to trade-offs. In addition, there could be biological limits to the possible combinations of vital rate estimates due to covariance between vital rates. While these factors could reduce the uncertainty impacts identified in this study, the exact magnitude of any such reduction remains uncertain.

## Data/code for peer review statement

All data and code used to produce the results for this study can be found at https://github.com/emilygsimmonds/parameter_uncertainty/tree/main. This will be updated to a Zenodo repository on acceptance after peer review.

## Acknowledgements

We would first like to thank Andreas Robert Dorf Nielsen and Alba Elisabeth Forum for their valuable help data checking our reviewed papers. Next, we would like to give thanks to Rob Salguero-Gómez for his helpful and constructive comments on this manuscript. We would also like to thank all those that have contributed to the construction, maintenance, and running of the COMADRE open access database. Without this resource research such as this would not be possible.

Emily G. Simmonds would like to acknowledge her funding for the PREDICT project from the Research Council of Norway: project number 314952.

## Author contributions

EGS led this project. She conceived of the initial idea, conducted the review, analysed results, and constructed and implemented the simulation study. EGS also led the writing of the manuscript.

ORJ contributed ideas on the conceptual development of this project, checked some review outputs, and has contributed substantially to the interpretation and writing up of the project results and final manuscript.

All authors have given their approval to the final manuscript and agree to be accountable for the aspects of the work they conducted.

## Statement on inclusion

Our study is based on a review of literature and a simulation study rather than primary data. As such, there was no local data collection and the data are theoretical and have no origin region. The authorship team in this case is very small but covers several European countries.

