## Supporting Information for "Uncertainty propagation in matrix population models: gaps, importance, and guidelines"

**Table of contents**

| S1: Estimating standard error from confidence intervals | 2 |
| --- | --- |
| S2: Propagation results for 2-by-2 and 5-by-5 matrices (mid-level uncertainty) | 3 |
| S3: Conclusion changes for 2-by-2 and 5-by-5 matrices (mid-level uncertainty) | 5 |
| S4: Elasticity changes for all matrix dimensions and uncertainty levels | 7 |
| S5: Propagation results for all matrix dimensions under low-level uncertainty | 10 |
| S6: Conclusion changes for all matrix dimensions under low-level uncertainty | 11 |
| S7: Propagation results for all matrix dimensions under high-level uncertainty | 14 |
| S8: Conclusion changes for all matrix dimensions under high-level uncertainty | 15 |
| S9: Table of reproduction:survival ratios | 18 |
| S10: Table of papers reviewed | 20 |

**S1: Estimating standard error from confidence intervals**

The process of estimating a standard error from a given confidence interval involved subtracting the vital rate estimate from the upper confidence interval and dividing the resulting difference by two. By carrying out this operation, we assumed that the reported confidence intervals were approximately symmetrical and corresponded to the scale of the estimate. To verify this assumption for each vital rate, we compared the reported lower confidence interval and the calculated value of the estimate minus twice the estimated standard error. If the reported confidence interval was within a margin of 0.05 (an allowance made to accommodate minor discrepancies and rounding differences) of the calculated lower confidence interval, we deemed that the assumptions of symmetry and scale were met. Confidence intervals that failed to satisfy this assumption were regarded as outliers and subsequently excluded from further analyses.

**S2: Propagation results for 2-by-2 and 5-by-5 matrices under mid-level uncertainty**

Below are the results of uncertainty in population growth rate under different propagation scenarios for matrix dimensions 2-by-2 and 5-by-5 under the median uncertainty level. Figures S1 and S2 show a similar pattern to Figure 2 in the main manuscript, with increasing importance of propagating fecundity uncertainty as the fecundity dominance of the matrix increases.

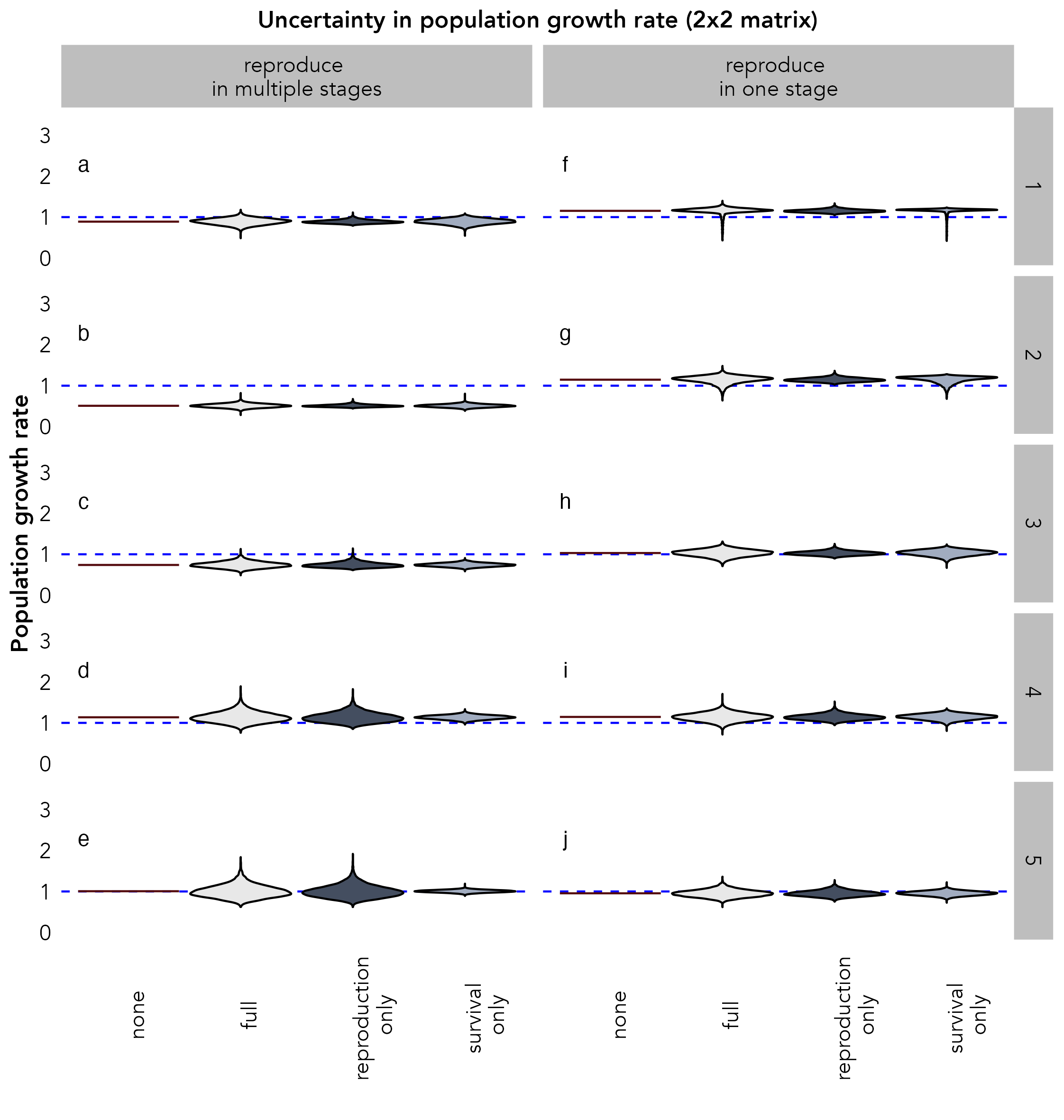

Figure S 1: The estimated population growth rate distribution under different types of uncertainty propagation and with different life histories, using parametric resampling. The left-hand column shows cases where reproduction occurs in multiple stages, while the right-hand column shows cases where reproduction occurs in a single stage only. The relative importance of fecundity vs survival changes systematically from rows 1 to 5 such that row 1 is relatively survival dominant, and row 5 is relatively fecundity dominant. The blue dashed line indicates a population growth rate of 1. All results are for a 2-by-2 matrix at the mid-uncertainty level.

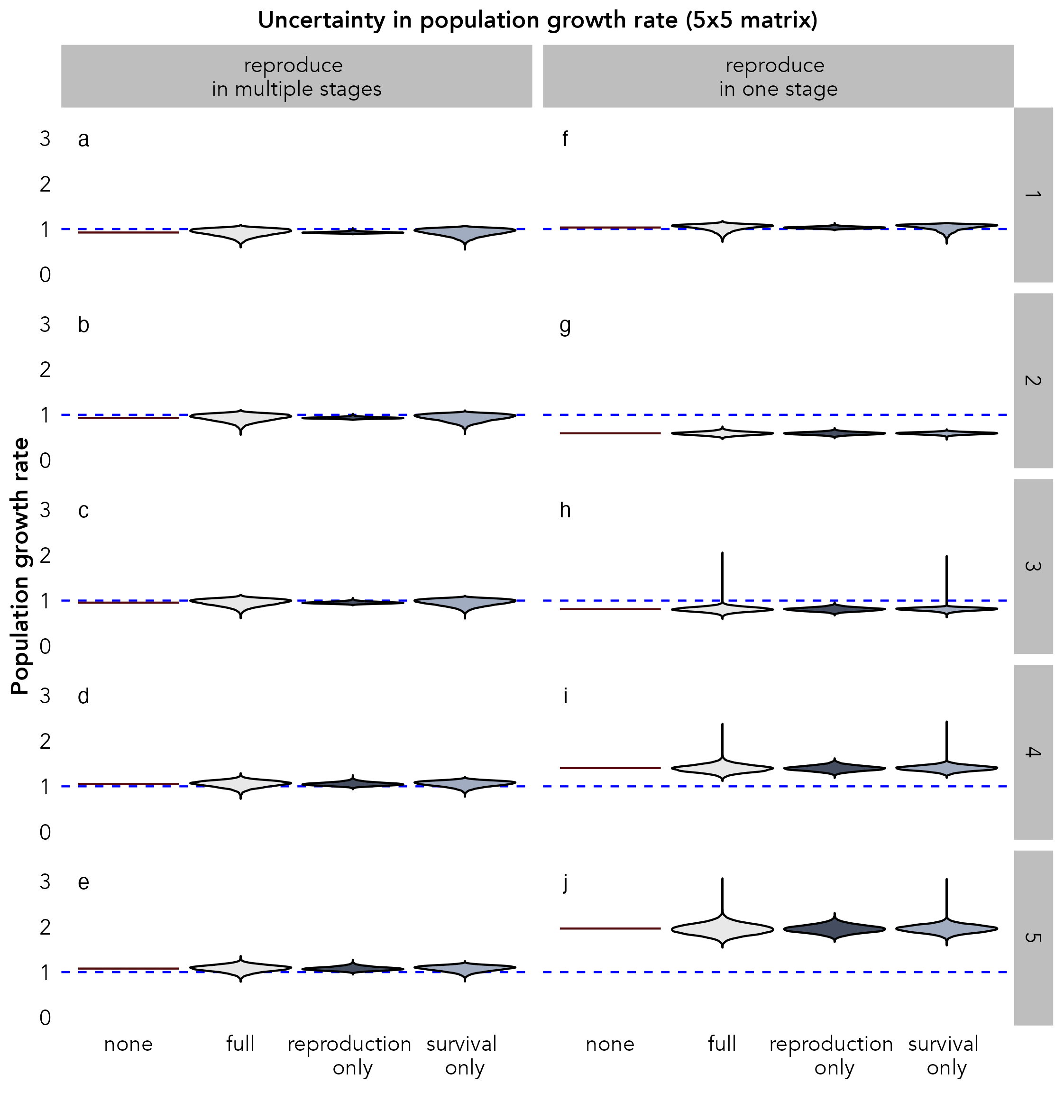

Figure S 2: The estimated population growth rate distribution under different types of uncertainty propagation and with different life histories, using parametric resampling. The left-hand column shows cases where reproduction occurs in multiple stages, while the right-hand column shows cases where reproduction occurs in a single stage only. The relative importance of fecundity vs survival changes systematically from rows 1 to 5 such that row 1 is relatively survival dominant, and row 5 is relatively fecundity dominant. The blue dashed line indicates a population growth rate of 1. All results are for a 5-by-5 matrix at the mid-uncertainty level.

**S3: Conclusion changes for 2-by-2 and 5-by-5 matrices under mid-level uncertainty**

Below are the results for the percentage of times the conclusion of the population trend changed during resampling for matrix dimensions 2-by-2 and 5-by-5 under the median uncertainty level. Figures S3 shows a similar pattern to Figure 3 in the main manuscript, however, Figure S4 highlights more clearly the bias that can be caused by partial propagation. The left-hand column of Figure S4 shows that only propagating fecundity uncertainty leads to an overconfident conclusion. In contrast, the right-hand column suggests species that reproduce only in one stage produced conclusions that were more robust, with little change under uncertainty propagation.

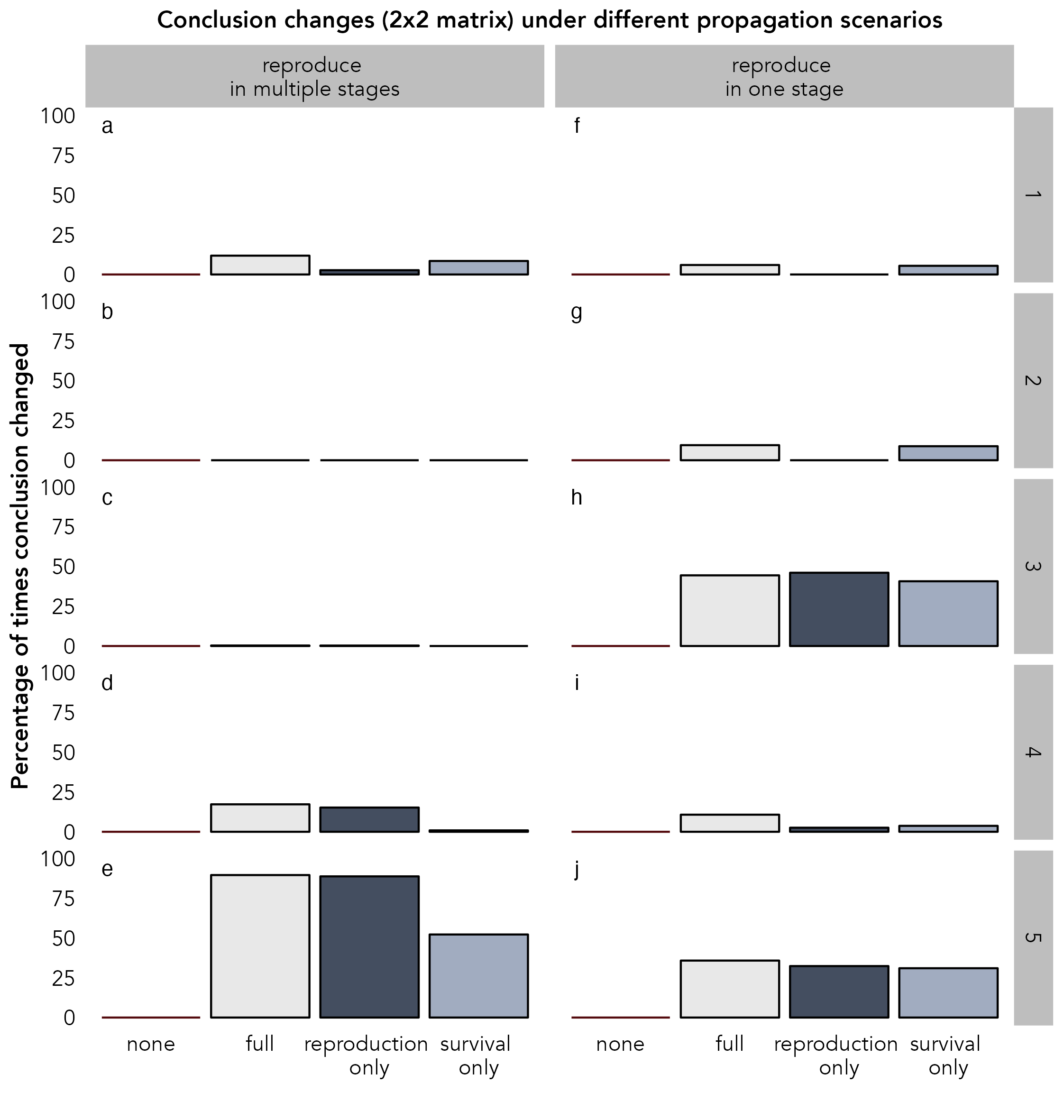

Figure S 3: The percentage of times a conclusion changed during parametric resampling under different types of uncertainty propagation, and with different life histories. The left-hand column shows cases where reproduction occurs in multiple stages, while the right-hand column shows cases where reproduction occurs in a single stage only. The relative importance of fecundity vs survival changes systematically from rows 1 to 5 such that row 1 is relatively survival dominant, and row 5 is relatively fecundity dominant. All results are for a 2-by-2 matrix at the mid-uncertainty level.

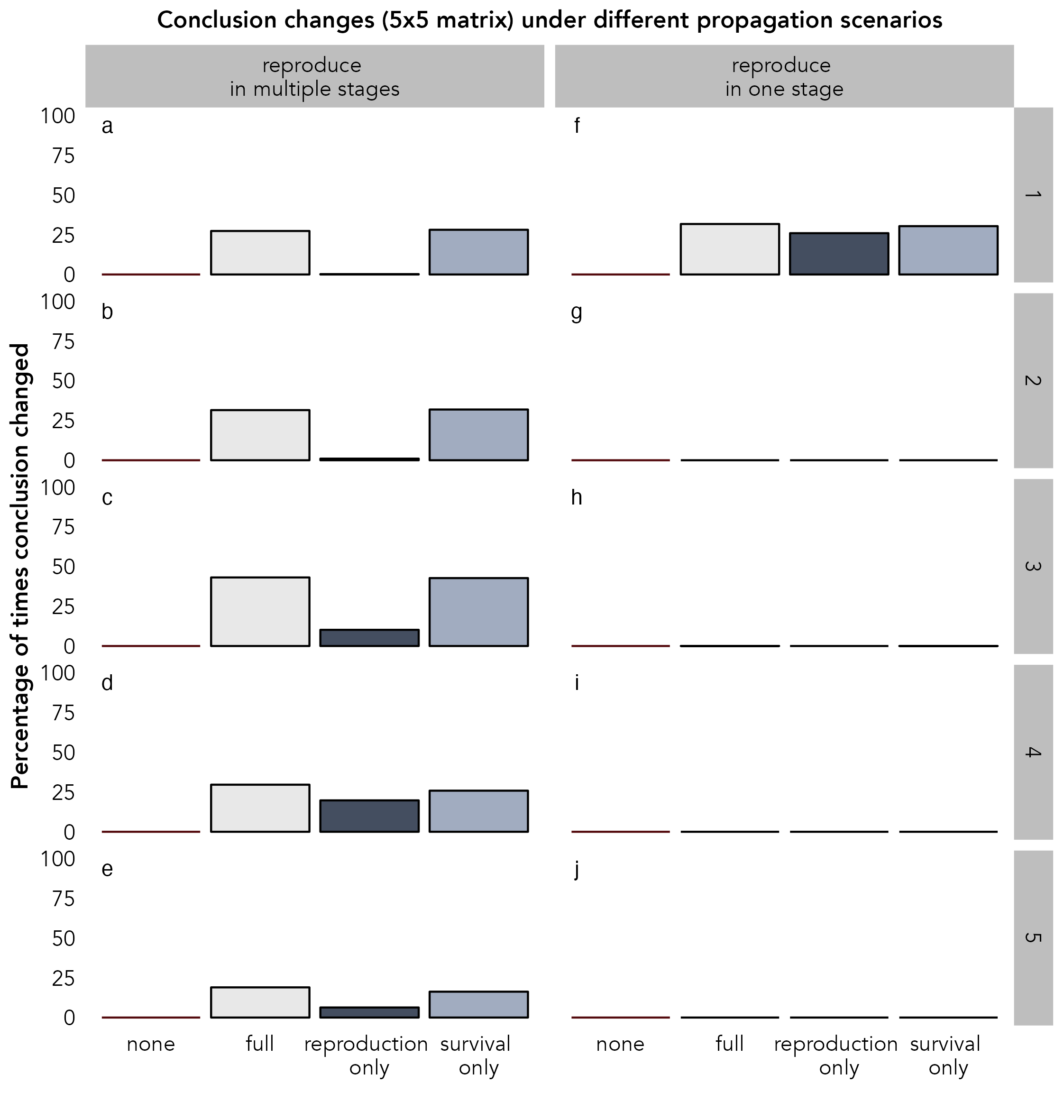

Figure S 4: The percentage of times a conclusion changed during parametric resampling under different types of uncertainty propagation, and with different life histories. The left-hand column shows cases where reproduction occurs in multiple stages, while the right-hand column shows cases where reproduction occurs in a single stage only. The relative importance of fecundity vs survival changes systematically from rows 1 to 5 such that row 1 is relatively survival dominant, and row 5 is relatively fecundity dominant. All results are for a 5-by-5 matrix at the mid-uncertainty level.

**S4: Elasticity changes for all matrix dimensions and uncertainty levels**

This section presents the percentage of times that the dominant vital rate (as identified through elasticities) changed during resampling. This is effectively an indication of how robust the identification of the dominant vital rate is to uncertainty propagation. Figures S5-7 shows that for a 2-by-2 matrix, the dominant vital rate does not change substantially even under high uncertainty (max. 50% of resamples). In contrast, there is more variability in the 3-by-3 matrix and 5-by-5 matrix results. While there is no change for most scenarios under low uncertainty, some life history combinations do produce a 25-100% change in the estimated dominant vital rate, even under low uncertainty. This suggests that even under low uncertainty for larger matrices the identification of dominant vital rates can be highly variable when uncertainty is accounted for. Under high uncertainty, all matrix dimensions and scenarios show some variation in the conclusions and caution should be applied when interpreting results of the dominant vital rate.

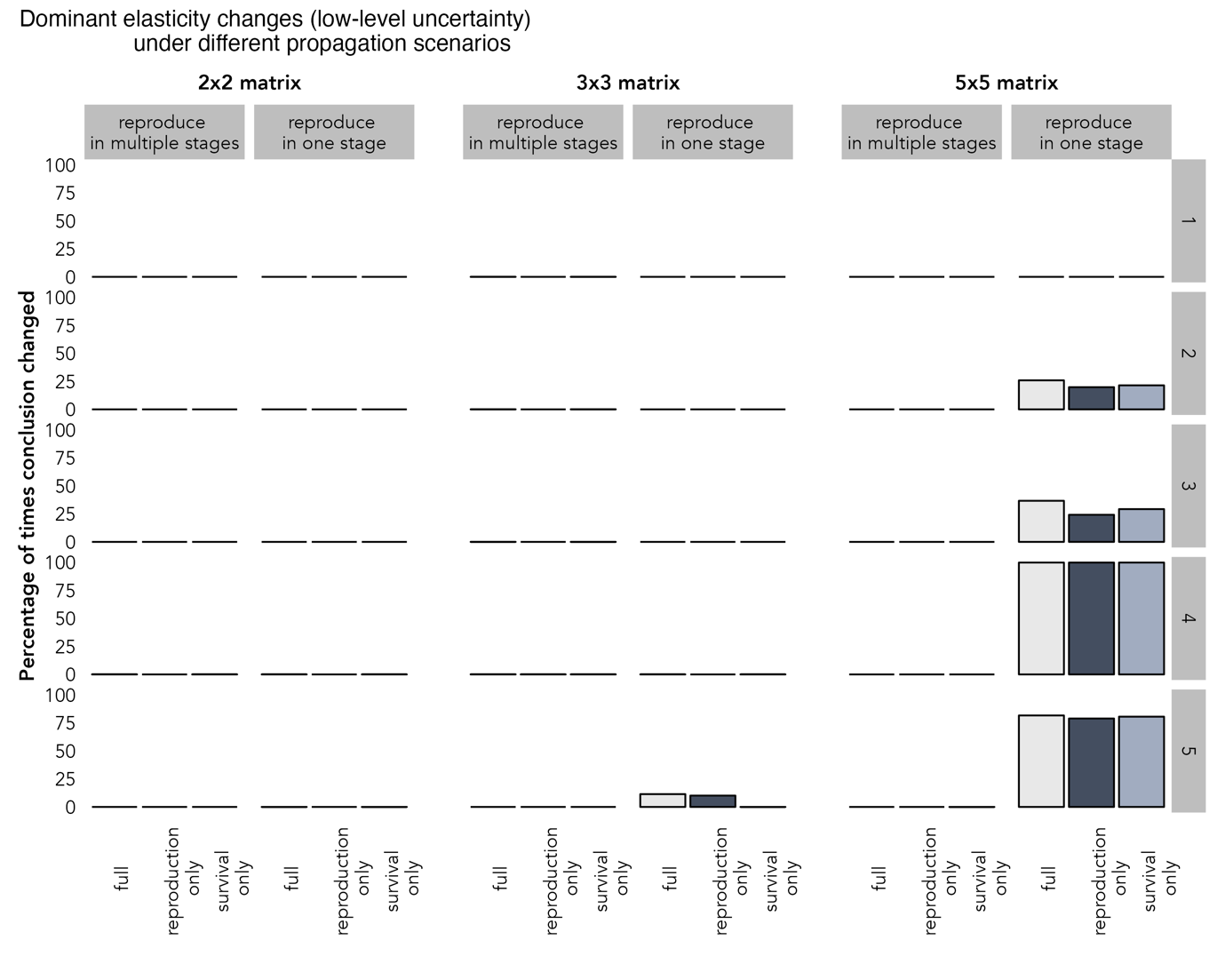

Figure S 5: The percentage of times the vital rate with the highest elasticity changed during parametric resampling under different types of uncertainty propagation, and with different life histories. The left-hand column of each panel shows cases where reproduction occurs in multiple stages, while the right-hand column shows cases where reproduction occurs in a single stage only. The relative importance of fecundity vs survival changes systematically from rows 1 to 5 such that row 1 is relatively survival dominant, and row 5 is relatively fecundity dominant. Results are for a low uncertainty level.

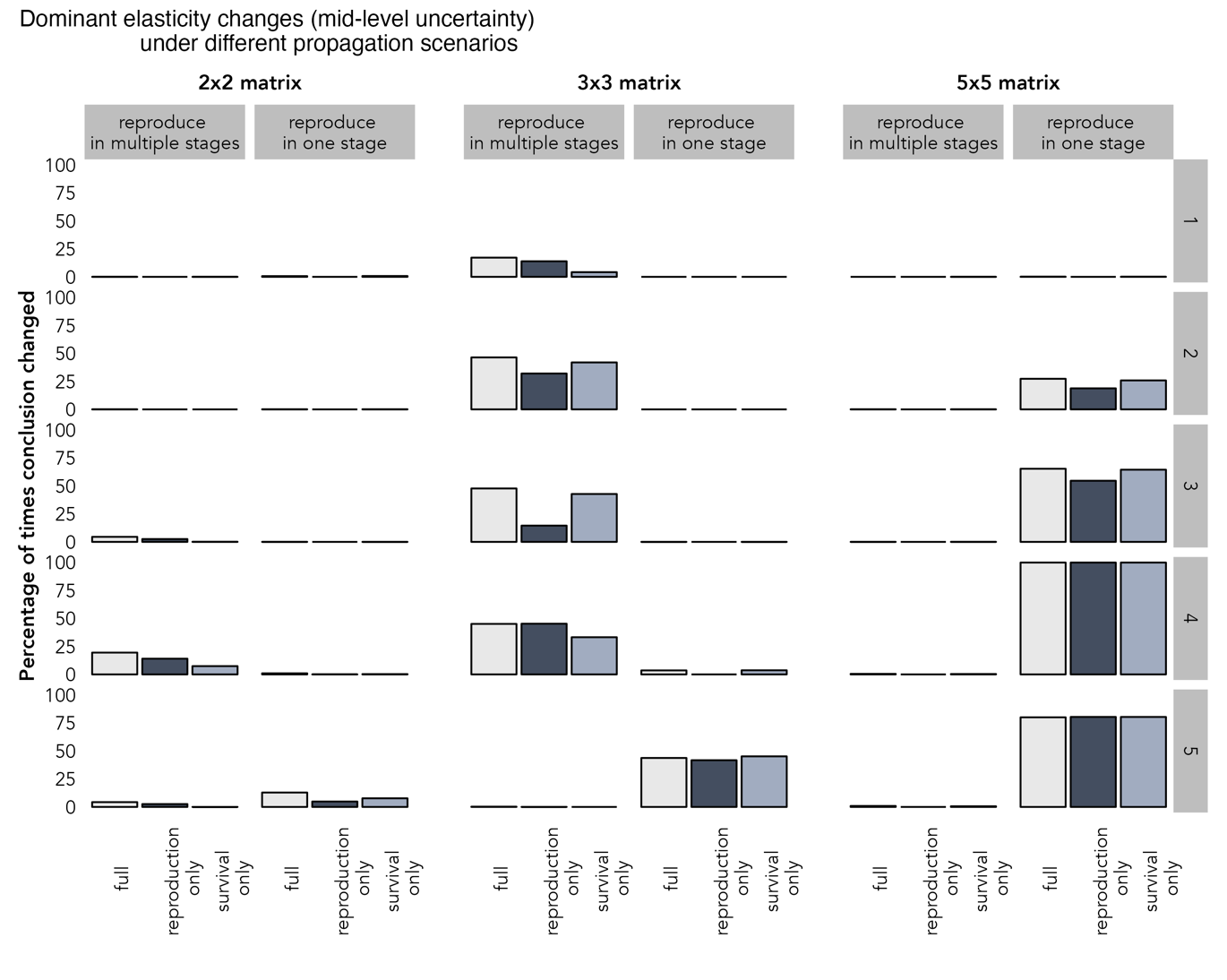

Figure S 6: The percentage of times the vital rate with the highest elasticity changed during parametric resampling under different types of uncertainty propagation, and with different life histories. The left-hand column of each panel shows cases where reproduction occurs in multiple stages, while the right-hand column shows cases where reproduction occurs in a single stage only. The relative importance of fecundity vs survival changes systematically from rows 1 to 5 such that row 1 is relatively survival dominant, and row 5 is relatively fecundity dominant. Results are for a mid uncertainty level.

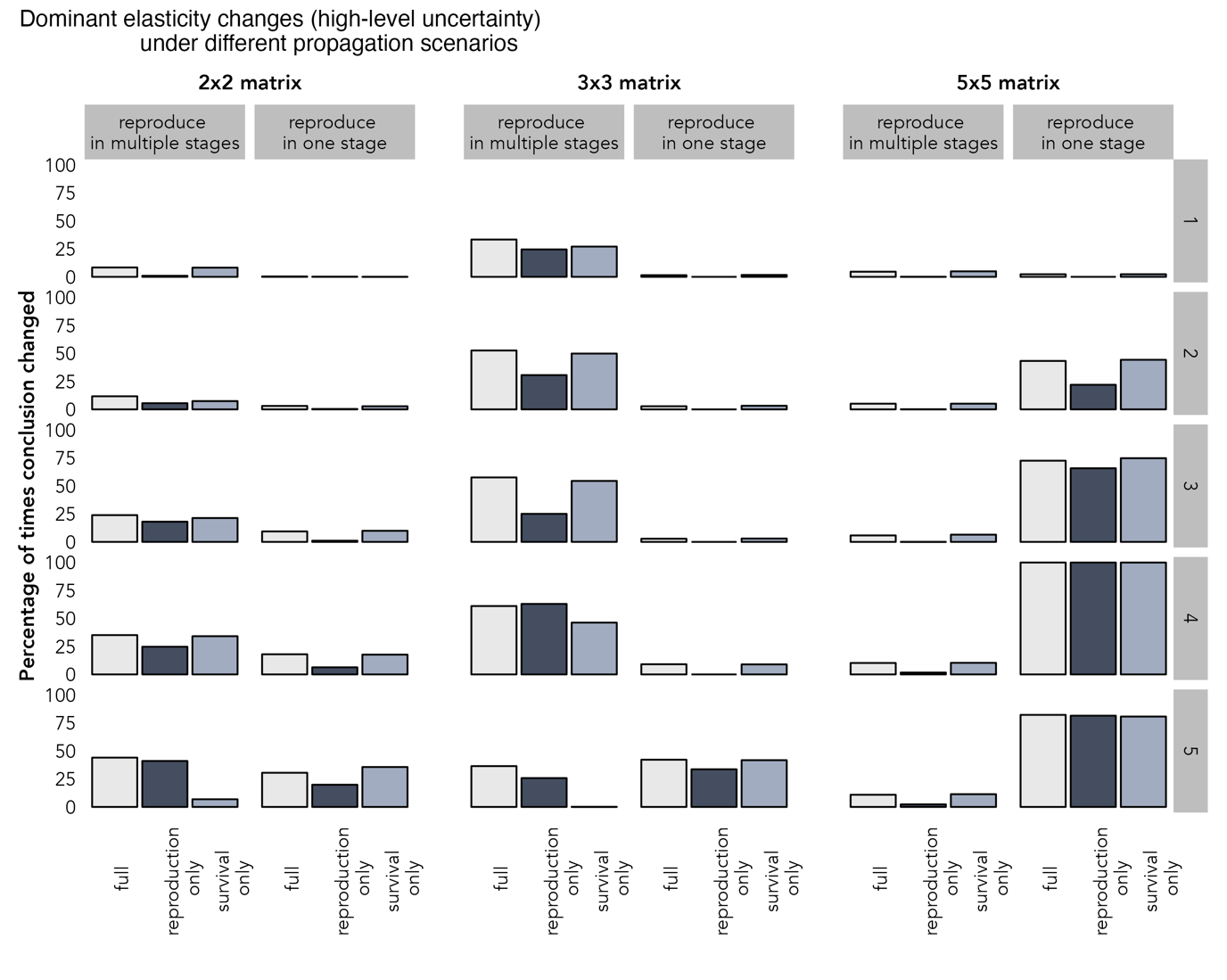

Figure S 7: The percentage of times the vital rate with the highest elasticity changed during parametric resampling under different types of uncertainty propagation, and with different life histories. The left-hand column of each ponel shows cases where reproduction occurs in multiple stages, while the right-hand column shows cases where reproduction occurs in a single stage only. The relative importance of fecundity vs survival changes systematically from rows 1 to 5 such that row 1 is relatively survival dominant, and row 5 is relatively fecundity dominant. Results are for a high uncertainty level.

**S5: Propagation results for all matrix dimensions under low-level uncertainty**

Below are the results of uncertainty in population growth rate under different propagation scenarios for all matrix dimensions under the low uncertainty level. Figure S8 generally shows much lower uncertainty distributions than Figure 2 in the main text.

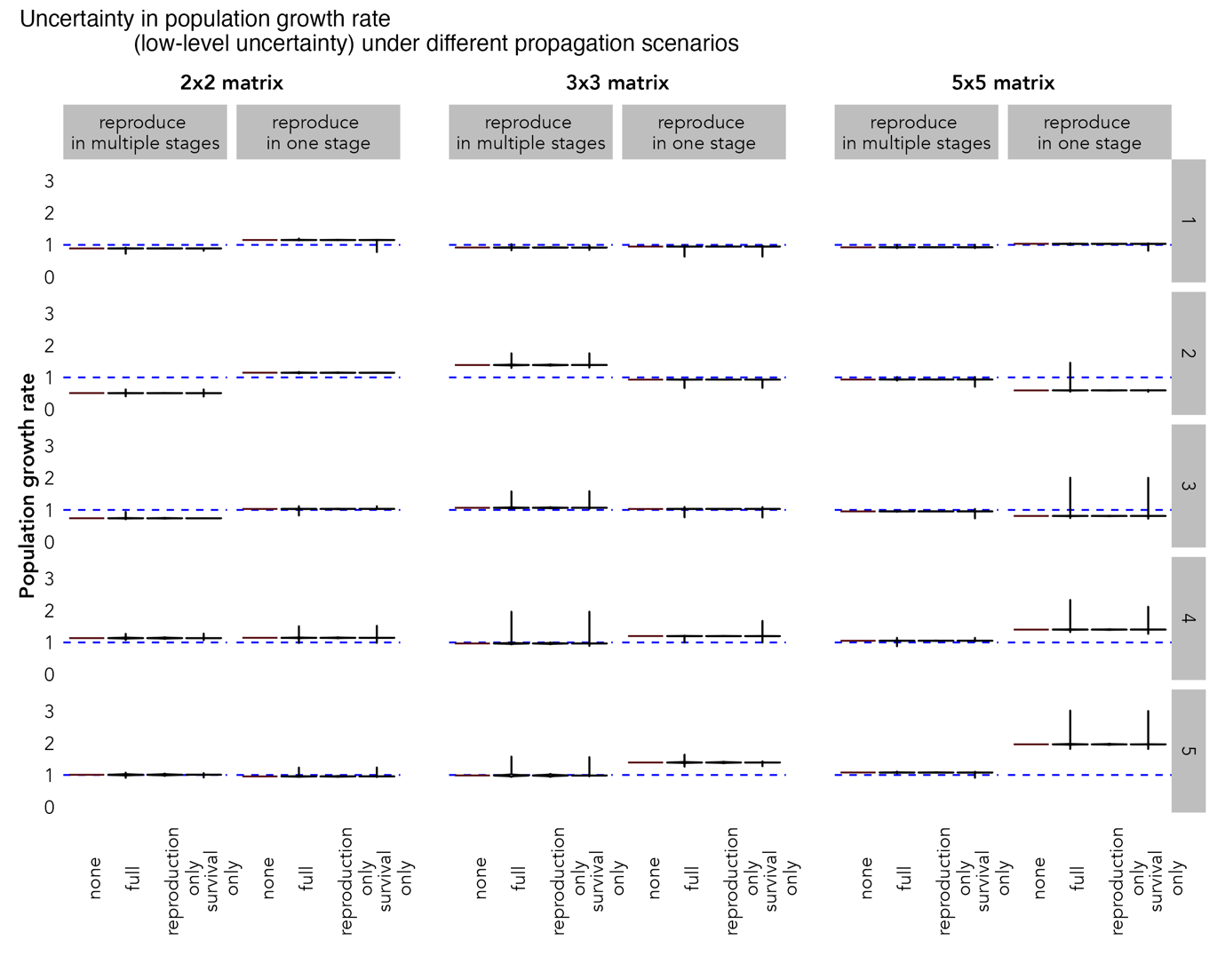

Figure S 8: The estimated population growth rate distribution under different types of uncertainty propagation and with different life histories, using parametric resampling. The left-hand column of each pair shows cases where reproduction occurs in multiple stages, while the right-hand column shows cases where reproduction occurs in a single stage only. The relative importance of fecundity vs survival changes systematically from rows 1 to 5 such that row 1 is relatively survival dominant, and row 5 is relatively fecundity dominant. The blue dashed line indicates a population growth rate of 1. All results are for the low-uncertainty level.

**S6: Conclusion changes for all matrix dimensions under low-level uncertainty**

Below are the results for the percentage of times the conclusion of the population trend changed during resampling for all matrix dimensions under the low uncertainty level. Figures S9-11 show almost no changes to conclusion under the lowest level of uncertainty we explored.

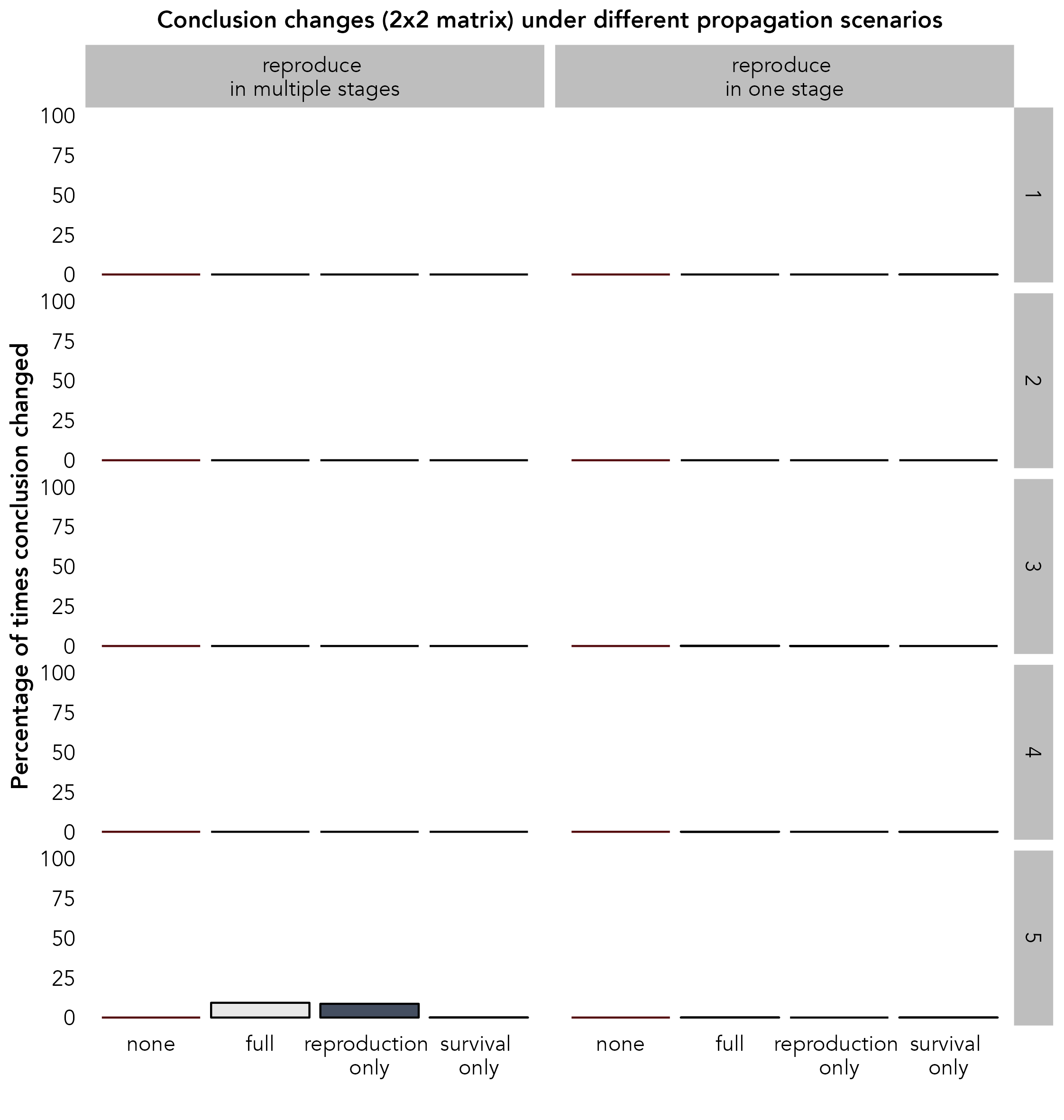

Figure S 9: The percentage of times a conclusion changed during parametric resampling under different types of uncertainty propagation, and with different life histories. The left-hand column shows cases where reproduction occurs in multiple stages, while the right-hand column shows cases where reproduction occurs in a single stage only. The relative importance of fecundity vs survival changes systematically from rows 1 to 5 such that row 1 is relatively survival dominant, and row 5 is relatively fecundity dominant. All results are for a 2-by-2 matrix at the low-uncertainty level.

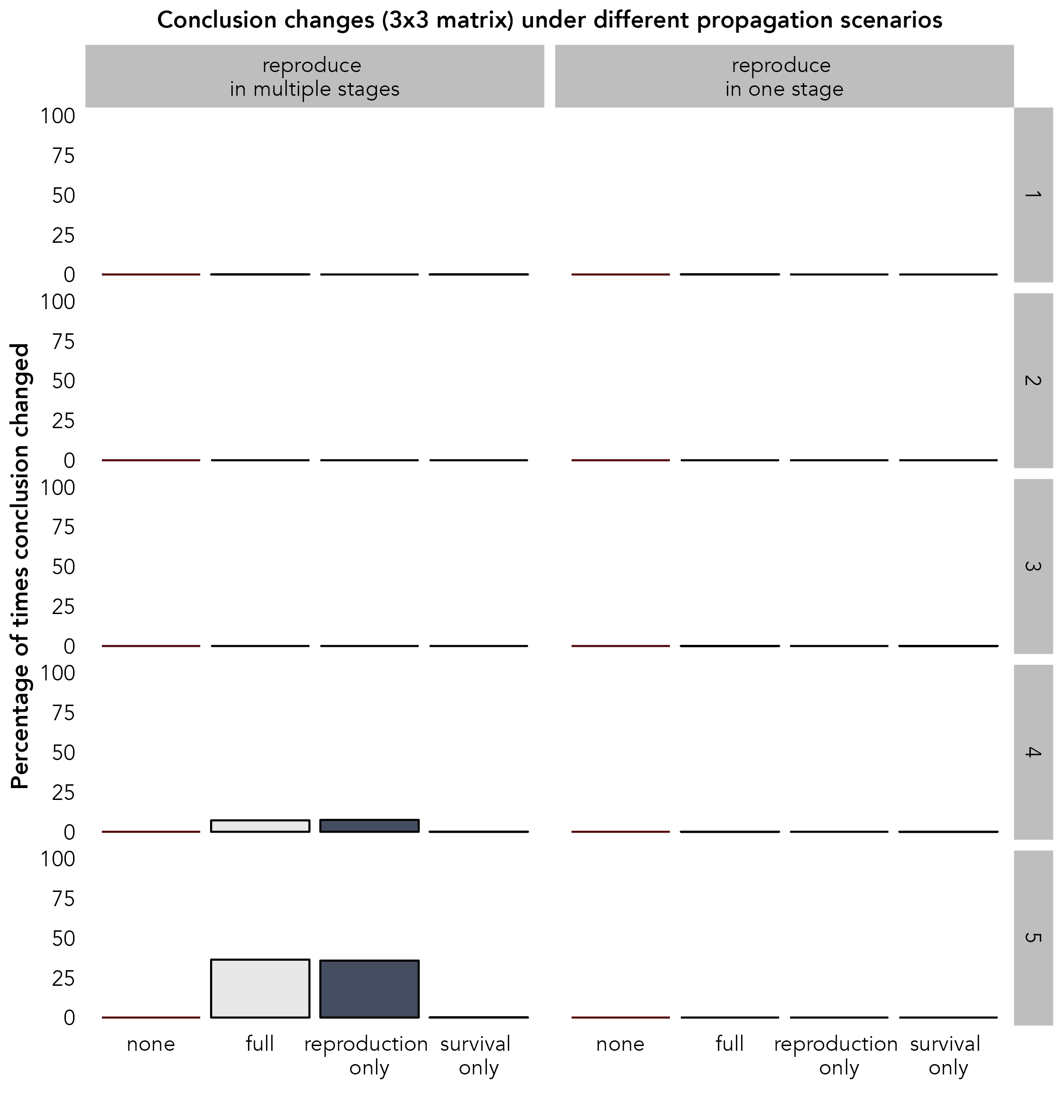

Figure S 10: The percentage of times a conclusion changed during parametric resampling under different types of uncertainty propagation, and with different life histories. The left-hand column shows cases where reproduction occurs in multiple stages, while the right-hand column shows cases where reproduction occurs in a single stage only. The relative importance of fecundity vs survival changes systematically from rows 1 to 5 such that row 1 is relatively survival dominant, and row 5 is relatively fecundity dominant. All results are for a 3-by-3 matrix at the low-uncertainty level.

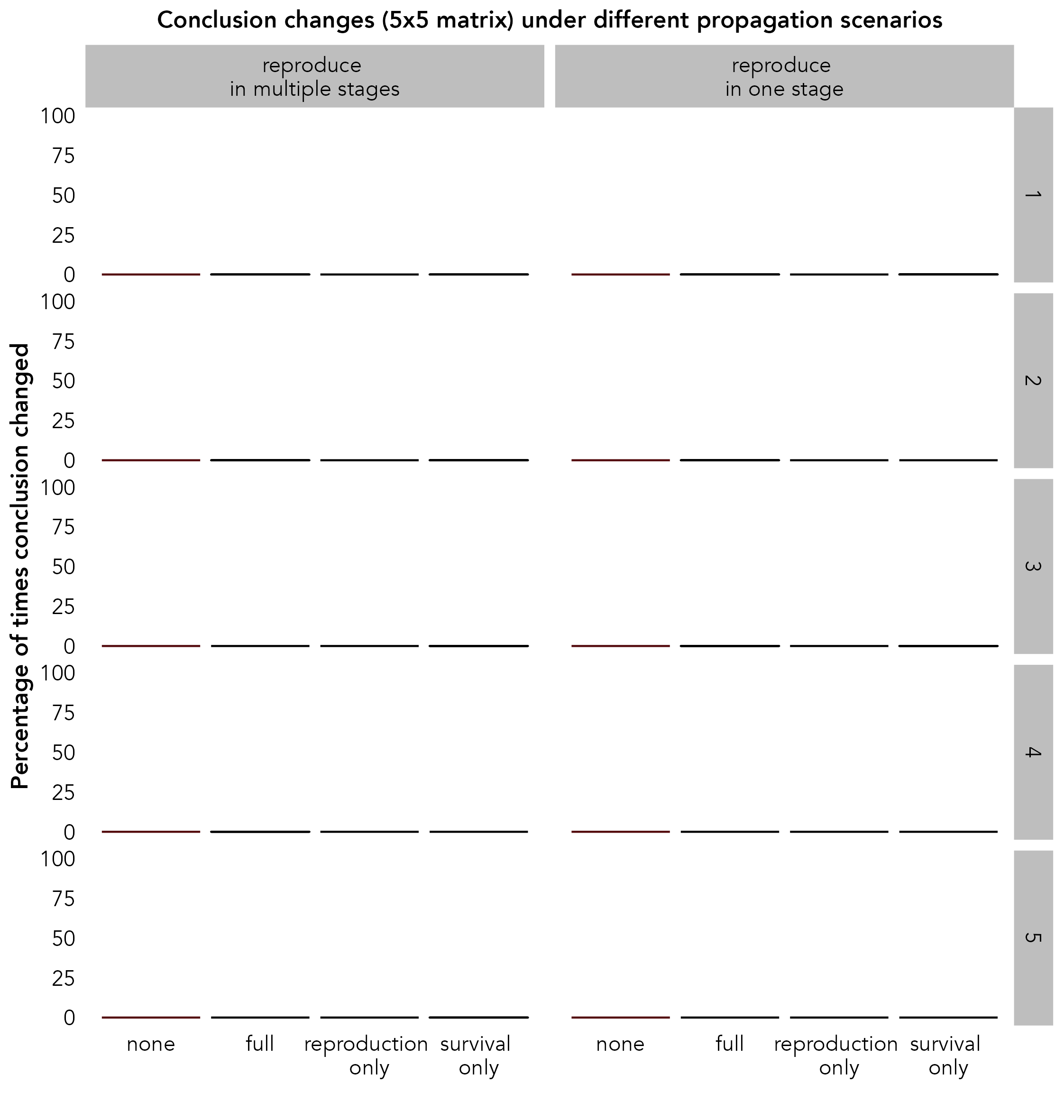

Figure S 11: The percentage of times a conclusion changed during parametric resampling under different types of uncertainty propagation, and with different life histories. The left-hand column shows cases where reproduction occurs in multiple stages, while the right-hand column shows cases where reproduction occurs in a single stage only. The relative importance of fecundity vs survival changes systematically from rows 1 to 5 such that row 1 is relatively survival dominant, and row 5 is relatively fecundity dominant. All results are for a 5-by-5 matrix at the low-uncertainty level.

**S7: Propagation results for all matrix dimensions under high-level uncertainty**

Below are the results of uncertainty in population growth rate under different propagation scenarios for all matrix dimensions under the high uncertainty level. Figure S12 generally shows much wider uncertainty distributions than Figure 2 in the main text. In particular, it should be noted the y-axis now extends to 10.

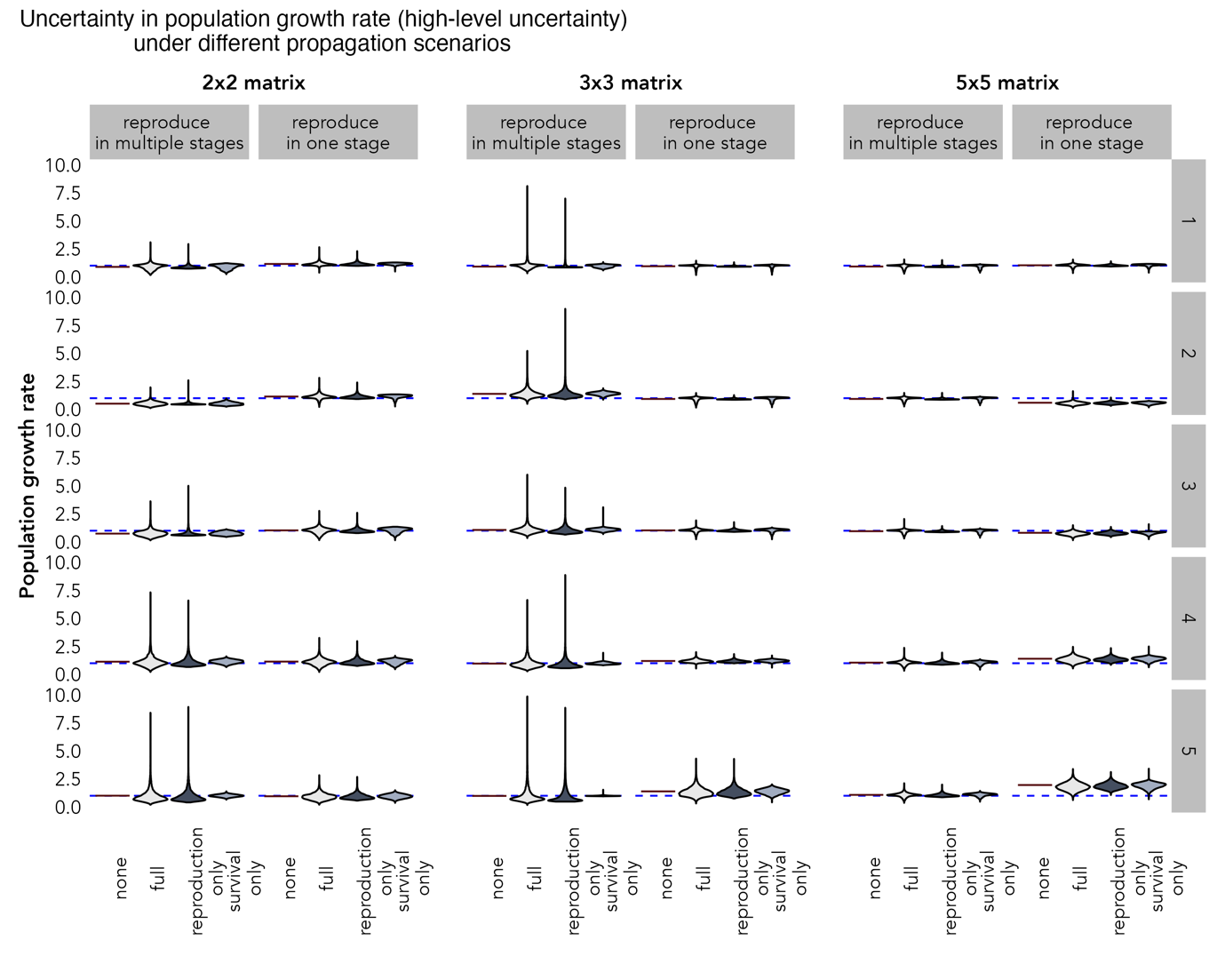

Figure S 12: The estimated population growth rate distribution under different types of uncertainty propagation and with different life histories, using parametric resampling. The left-hand column of each pair shows cases where reproduction occurs in multiple stages, while the right-hand column shows cases where reproduction occurs in a single stage only. The relative importance of fecundity vs survival changes systematically from rows 1 to 5 such that row 1 is relatively survival dominant, and row 5 is relatively fecundity dominant. The blue dashed line indicates a population growth rate of 1. All results are for the high-uncertainty level. Some values were not plotted as they fell outside of the bounds of plot (n = 2 for the 2-by-2 matrices and n = 7 for the 3-by-3 matrices). Extending the plotting axes to include these points made viewing of the bulk of the distribution challenging therefore we opted to omit them to improve clarity.

**S8: Conclusion changes for all matrix dimensions under low-level uncertainty**

Below are the results for the percentage of times the conclusion of the population trend changed during resampling for all matrix dimensions under the high uncertainty level. Figures S13-15 show that substantial conclusion changes are present in most cases we explored under the highest uncertainty level. The bias caused by partial propagation is still highlighted, especially for species that reproduce in multiple stages.

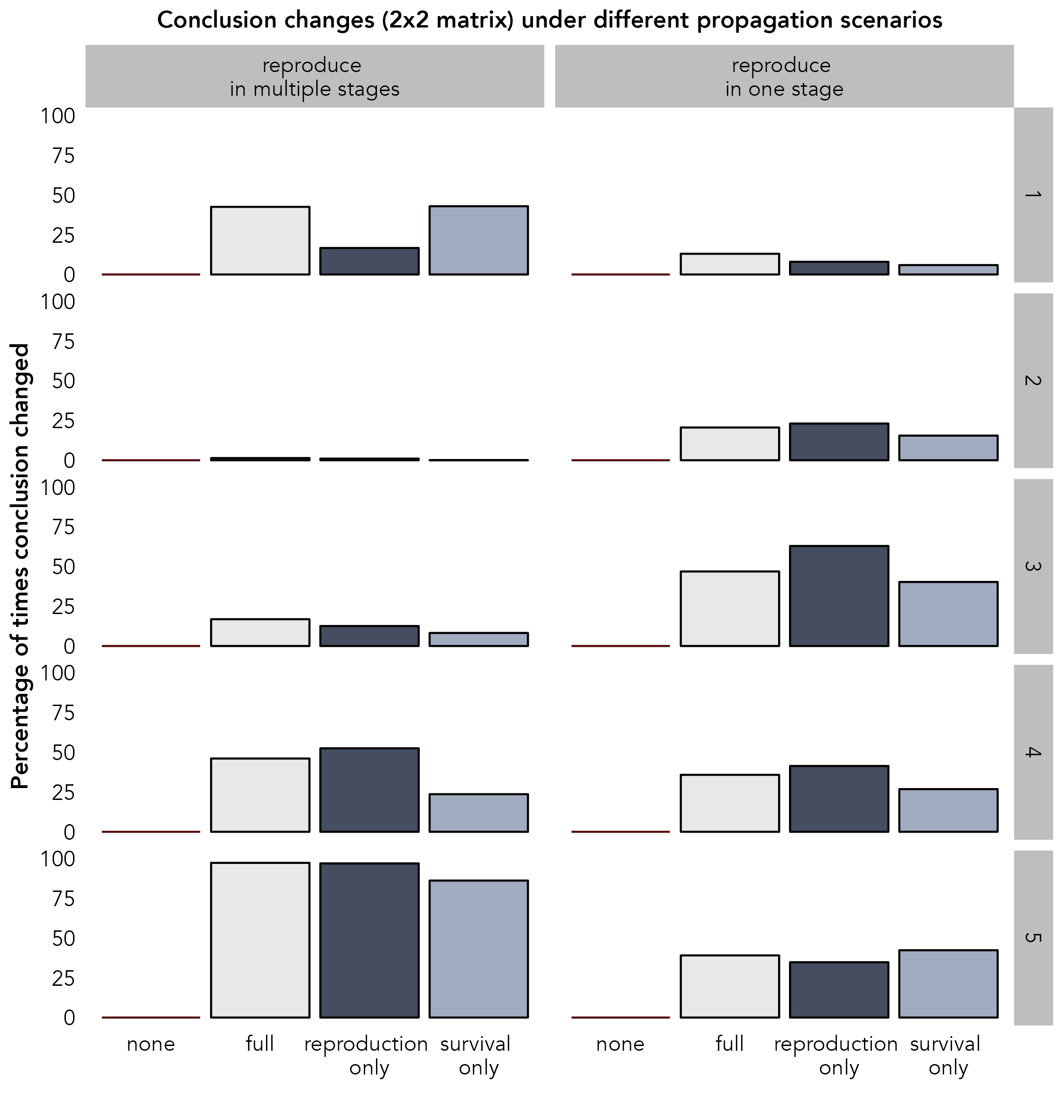

Figure S 13: The percentage of times a conclusion changed during parametric resampling under different types of uncertainty propagation, and with different life histories. The left-hand column shows cases where reproduction occurs in multiple stages, while the right-hand column shows cases where reproduction occurs in a single stage only. The relative importance of fecundity vs survival changes systematically from rows 1 to 5 such that row 1 is relatively survival dominant, and row 5 is relatively fecundity dominant. All results are for a 2-by-2 matrix at the high-uncertainty level.

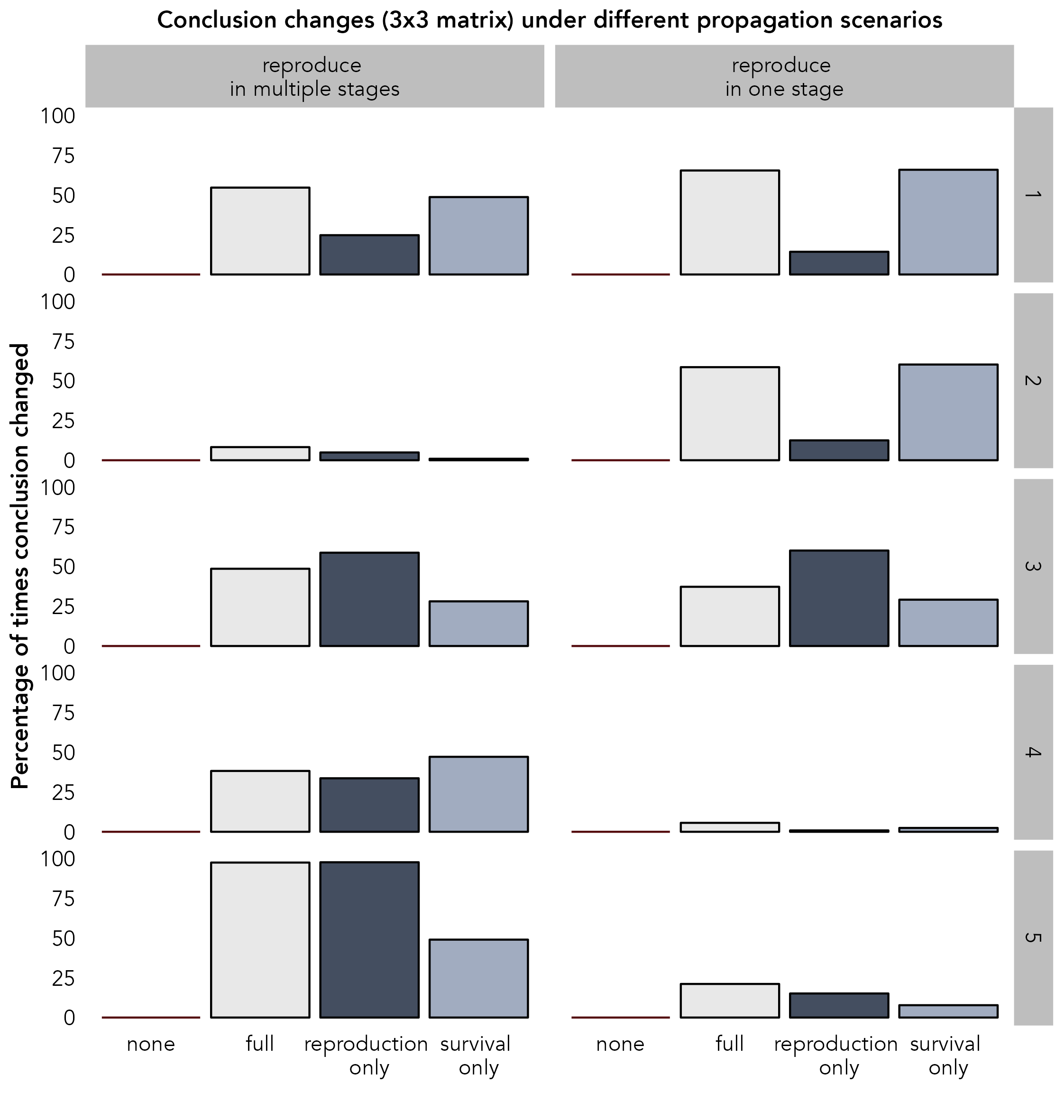

Figure S 14: The percentage of times a conclusion changed during parametric resampling under different types of uncertainty propagation, and with different life histories. The left-hand column shows cases where reproduction occurs in multiple stages, while the right-hand column shows cases where reproduction occurs in a single stage only. The relative importance of fecundity vs survival changes systematically from rows 1 to 5 such that row 1 is relatively survival dominant, and row 5 is relatively fecundity dominant. All results are for a 3-by-3 matrix at the high-uncertainty level.

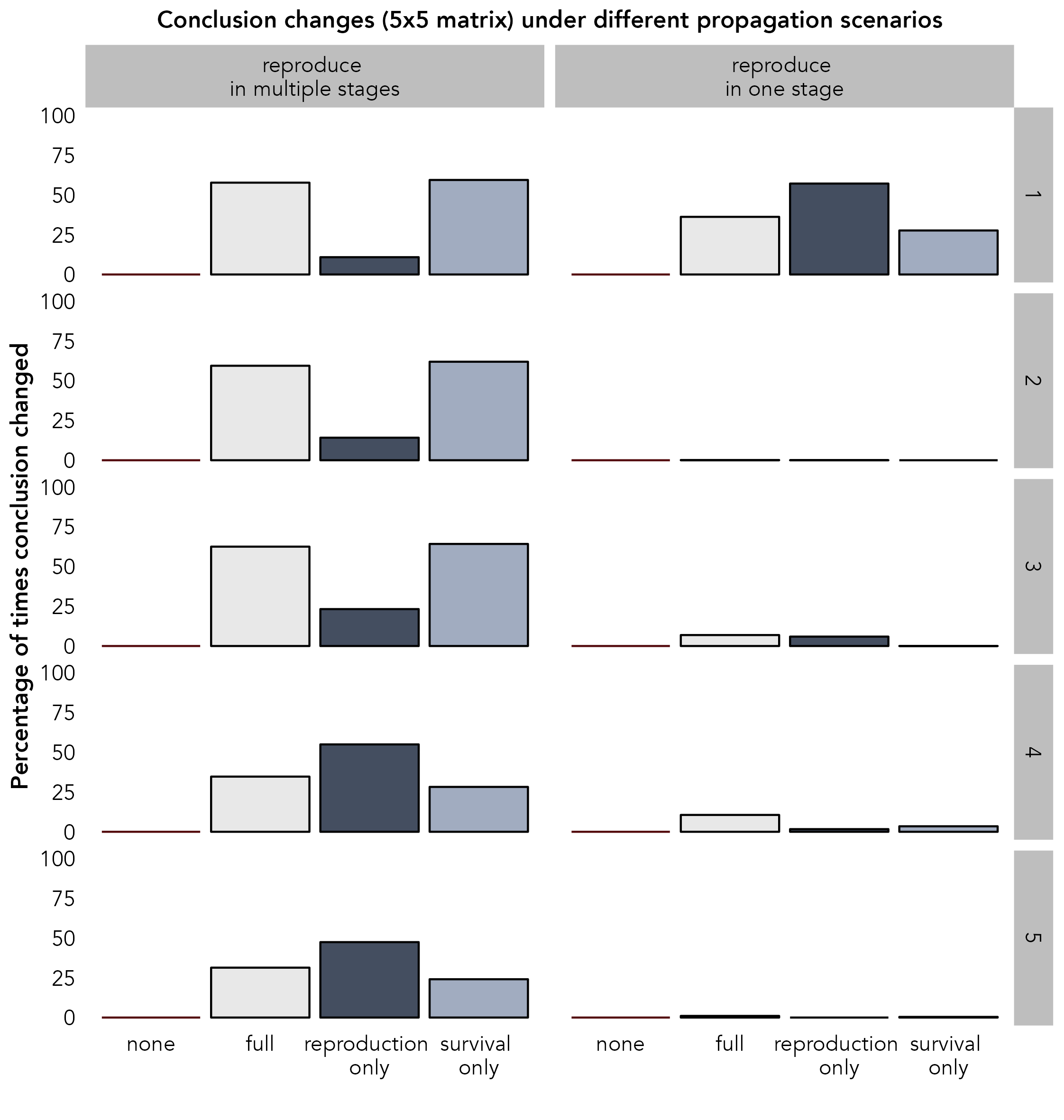

Figure S 15: The percentage of times a conclusion changed during parametric resampling under different types of uncertainty propagation, and with different life histories. The left-hand column shows cases where reproduction occurs in multiple stages, while the right-hand column shows cases where reproduction occurs in a single stage only. The relative importance of fecundity vs survival changes systematically from rows 1 to 5 such that row 1 is relatively survival dominant, and row 5 is relatively fecundity dominant. All results are for a 5-by-5 matrix at the high-uncertainty level.

**S9: Table of reproduction:survival ratios**

Table S 1:Table of reproduction:survival ratios for all selected base matrices. Note that for some breeding strategies, all matrices are survival dominant (ratio of <1).

| Matrix number | Reproduction:Survival ratio | Reproduction strategy | Matrix size | Generation  Time (average age difference between parents and offspring) |
| --- | --- | --- | --- | --- |
| 1 | 0.22881267 | multiple | 2-by-2 | 5.37 |
| 2 | 0.28675204 | multiple | 2-by-2 | 4.49 |
| 3 | 0.4978409 | multiple | 2-by-2 | 3.01 |
| 4 | 0.78884475 | multiple | 2-by-2 | 2.27 |
| 5 | 1.62668407 | multiple | 2-by-2 | 1.61 |
| 1 | 0.31732463 | once | 2-by-2 | 7.3 |
| 2 | 0.41281293 | once | 2-by-2 | 5.84 |
| 3 | 0.50597812 | once | 2-by-2 | 4.95 |
| 4 | 0.70762381 | once | 2-by-2 | 3.83 |
| 5 | 0.87565845 | once | 2-by-2 | 3.28 |
| 1 | 0.45026608 | multiple | 3-by-3 | 4.7 |
| 2 | 0.68104032 | multiple | 3-by-3 | 3.45 |
| 3 | 1.01550905 | multiple | 3-by-3 | 2.97 |
| 4 | 1.6042943 | multiple | 3-by-3 | 2.04 |
| 5 | 9.29498546 | multiple | 3-by-3 | 1.18 |
| 1 | 0.17064779 | once | 3-by-3 | 18.58 |
| 2 | 0.23346264 | once | 3-by-3 | 13.85 |
| 3 | 0.35883266 | once | 3-by-3 | 9.36 |
| 4 | 0.52210945 | once | 3-by-3 | 8.66 |
| 5 | 0.99311912 | once | 3-by-3 | 3.01 |
| 1 | 0.13033525 | multiple | 5-by-5 | 14.43 |
| 2 | 0.14520726 | multiple | 5-by-5 | 13.05 |
| 3 | 0.17771024 | multiple | 5-by-5 | 10.85 |
| 4 | 0.30231316 | multiple | 5-by-5 | 6.79 |
| 5 | 0.33009495 | multiple | 5-by-5 | 6.3 |
| 1 | 0.43244405 | once | 5-by-5 | 12.56 |
| 2 | 1 | once | 5-by-5 | 5 |
| 3 | 1 | once | 5-by-5 | 1.32e-15 |
| 4 | 1.23255364 | once | 5-by-5 | 5.87 |
| 5 | 1.32609582 | once | 5-by-5 | 5.52 |

While the reproduction:survival ratio is designed primarily to look at which vital rates contribute most to changes in population growth rate ($\lambda$), it also shows a correlation with generation time (average age difference between parents and offspring) for most size by reproductive strategy groups. Generation time was calculated using the gen_time() function in the Rage R package (Jones, Barks, Stott, et al., 2022). As the ratio increases the generation time decreases, suggesting a move to a faster life history strategy as reproductive processes have greater influence on $\lambda$. This ratio-generation time correlation occurs for all groups except the five-by-five matrices for species that breed only in one stage. For this latter group, there is no discernible relationship between generation time and reproduction:survival ratio, therefore these matrices do not represent a clear life history strategy gradient.

**S10: Table of papers reviewed**

Table S 2: Details of all papers selected for review and reasons for any exclusion.

| Authors | Journal | Year | DOI_ISBN | Reason for  exclusion |
| --- | --- | --- | --- | --- |
| Hernandez-Pacheco; Rawlins; Kessler; Williams; Ruiz-Maldonado; Gonzalez-Martinez; Ruiz-Lambides; Sabat | Am J Primatol | 2013 | 10.1002/ajp.22177 |  |
| Kessler; Pacheco; Rawlings; Ruiz-Lambrides; Delgado; Sabat | Am J Primatol | 2014 | 10.1002/ajp.22323 |  |
| Hernandez-Pacheco; Delgado; Rawlins; Kessler; Ruiz-Lambides; Maldonado; Sabat | Am J Primatol | 2015 | 10.1002/ajp.22375 |  |
| Foster; Vincent | Aquat Conserv | 2012 | 10.1002/aqc.2243 |  |
| Davis; Hooten; Phillips; Doherty | Ecol Evol | 2014 | 10.1002/ece3.1290 |  |
| Gruebler; Korner-Nievergelt; Naef-Daenzer | Ecol Evol | 2014 | 10.1002/ece3.984 |  |
| Hinke; Trivelpiece; Trivelpiece | Ecosphere | 2017 | 10.1002/ecs2.1666 |  |
| Miller; Tietge; McMaster; Munkittrick; Xia; Griesmer; Ankley | Environ Toxicol | 2015 | 10.1002/etc.2972 |  |
| Reynolds; Weiser; Jamieson; Hatfield | J Wildlife Manage | 2013 | 10.1002/jwmg.582 |  |
| Chastant; King; Weseloh; Moore | J Wildlife Manage | 2014 | 10.1002/jwmg.628 |  |
| Chitwood; Lashley; Kilgo; Moorman; Deperno | J Wildlife Manage | 2015 | 10.1002/jwmg.835 |  |
| Edmunds | Limnol Oceanogr | 2015 | 10.1002/lno.10075 |  |
| Howerter; Anderson; Devries; Joynt; Armstrong; Emery; Arnold | Wildlife Monogr | 2014 | 10.1002/wmon.1012 |  |
| Haslob; Hauss; Petereit; Clemmesen; Kraus; Peck | Mar Biol | 2012 | 10.1007/s00227-012-1933-6 |  |
| Meyer; Robertson; Chilvers; Krkosek | Mar Biol | 2015 | 10.1007/s00227-015-2695-8 |  |
| Mercado-Molina; Ruiz-Diaz; Perez; Rodriguez-Barreras; Sabat | Coral Reefs | 2015 | 10.1007/s00338-015-1341-8) |  |
| Hudgens; Garcelon | Oecologia | 2011 | 10.1007/s00442-010-1761-7 |  |
| Grey | Oecologia | 2011 | 10.1007/s00442-011-1931-2 |  |
| Schaub; Reichlin; Abadi; Kery; Jenni; Arlettaz | Oecologia | 2012 | 10.1007/s00442-011-2070-5 |  |
| Ålvarez; Fernandez-Chacon; Genovart; Cano; Ojanguren; Rodriguez-Munoz; Nicieza | Oecologia | 2015 | 10.1007/s00442-015-3222-9 | No MPM |
| Hyslop; Stevenson; Macey; Carlile; Jenkins; Hostetler; Oli | Popul Ecol | 2011 | 10.1007/s10144-011-0292-3 |  |
| Kerbiriou; Le Viol; Bonnet; Robert | Popul Ecol | 2012 | 10.1007/s10144-012-0306-9 |  |
| Wolf; Hellgren; Schauber; Bogosian III; Kazmaier; Ruthven III; Moody | Popul Ecol | 2014 | 10.1007/s10144-014-0450-5 |  |
| Strauss; Kilewo; Rentsch; Packer | Popul Ecol | 2015 | 10.1007/s10144-015-0499-9 |  |
| Grear | Popul Ecol | 2016 | 10.1007/s10144-016-0562-1 |  |
| Nicol-Harper; Dooley; Packman; Mueller; Bijak; Hodgson; Townley; Ezard | Popul Ecol | 2018 | 10.1007/s10144-018-0620-y |  |
| Szostek | J Ornithol | 2011 | 10.1007/s10336-011-0745-7 |  |
| Duckworth; Altwegg; Harebottle | J Ornithol | 2011 | 10.1007/s10336-011-0758-2 |  |
| Duchet; Coutellec; Franquet; Lagneau; Lagadic | Ecotoxicology | 2010 | 10.1007/s10646-010-0507-y |  |
| Blomquist; Sade; Berard | Int J Primatol | 2010 | 10.1007/s10764-010-9461-z |  |
| Hovick; Miller | Landscape Ecol | 2013 | 10.1007/s10980-013-9896-7 | No MPM |
| Makenov; Bekova | Urban Ecosyst | 2016 | 10.1007/s11252-016-0566-9 |  |
| Santadino; Coviella; Momo | Water Air Soil Poll | 2014 | 10.1007/s11270-014-2207-3 |  |
| Sergio; Tavecchia; Blas; Lopez; Tanferna; Hiraldo | Basic Appl Ecol | 2011 | 10.1016/j.baae.2010.11.004 |  |
| Kruger; Grunkorn; Struwe-Juhl | Biol Conserv | 2010 | 10.1016/j.biocon.2009.12.010 |  |
| Rhodes; Ng; de Villiers; Preece; McAlpine; Possingham | Biol Conserv | 2011 | 10.1016/j.biocon.2010.12.027 |  |
| Chelliah; Bukka; Sukumar | Biol Conserv | 2013 | 10.1016/j.biocon.2013.05.008 |  |
| Wielgus; Morrison; Cooley; Maletzke | Biol Conserv | 2013 | 10.1016/j.biocon.2013.07.008 |  |
| Cruz; Pech; Seddon; Cleland; Nelson; Sanders; Maloney | Biol Conserv | 2013 | 10.1016/j.biocon.2013.09.005 |  |
| Goswami; Vasudev; Oli | Biol Conserv | 2014 | 10.1016/j.biocon.2014.05.026 |  |
| Walker; Marzluff; Cimprich | Biol Conserv | 2016 | 10.1016/j.biocon.2016.09.016. |  |
| Bergek; Ma; Vetemaa; Franz√©n; Appelberg | Ecotox Environ Safe | 2012 | 10.1016/j.ecoenv.2012.01.019 |  |
| Wiederholt; Fernandez-Duque; Diefenbach; Rudran | Ecol Model | 2010 | 10.1016/j.ecolmodel.2010.06.026 | No MPM |
| Lewis; Breck; Wilson; Webb | Ecol Model | 2014 | 10.1016/j.ecolmodel.2014.08.021 |  |
| Otjacques; De Laender; Kestemont | Ecol Model | 2016 | 10.1016/j.ecolmodel.2015.12.002 |  |
| Oli; Loughry; Caswell; Perez-Heydrich; McDonough; Truman | Ecol Model | 2017 | 10.1016/j.ecolmodel.2017.02.001 |  |
| Maestri; Ferrati; Berkunsky | Ecol Model | 2017 | 10.1016/j.ecolmodel.2017.07.023 |  |
| Rodriguez-Barreras; Perez; Mercado-Molina; Sabat | Estuar Coast Shelf S | 2015 | 10.1016/j.ecss.2015.06.021 |  |
| Stratford; Pollino; Brown | Environ Modell Softw | 2016 | 10.1016/j.envsoft.2016.02.009 |  |
| Mercado-Molina; Sabat; Yoshioka | J Exp Mar Biol Ecol | 2011 | 10.1016/j.jembe.2011.07.018 |  |
| Ferreira; Kajin; Vieira; Zangrandi; Cerqueira; Gentile | Mamm Biol | 2013 | 10.1016/j.mambio.2013.03.002 | Couldn't access: paywall |
| Pickett; Chan; Cheng; Allcock; Chan; Hu; Lee; Smith; Xing; Yu; Bonebrake | Environ Conserv | 2017 | 10.1017/S0376892917000340 |  |
| Hayman; McCrea; Restif; Suu-Ire; Fooks; Wood; Cunningham; Rowcliffe | J Mammal | 2012 | 10.1017/S0950268812000167 | No MPM |
| Van de Walle; Pigeon; Zedrosser; Swenson; Pelletier | Nature | 2018 | 10.1038/s41467-018-03506-3 |  |
| Chambers; Bencini | Wildlife Res | 2010 | 10.1071/WR10080 |  |
| Ng; Fredericks; Quist | N Am J Fish Manage | 2016 | 10.1080/02755947.2015.1111279 |  |
| Morris; Altmann; Brockman; Cords; Fedigan; Pusey; Stoinski; Bronikowski; Alberts; Strier | Am Nat | 2011 | 10.1086/657443 |  |
| Tsai; Sun; Punt; Liu | ICES J Mar Sci | 2014 | 10.1093/icesjms/fsu056 |  |
| Hanly; Haase | J Med Entomol | 2016 | 10.1093/jme/tjw021 |  |
| Henschke; Smith; Everett; Suthers | J Planck Res | 2015 | 10.1093/plankt/fbv024 |  |
| Edeline; Haugen; Weltzien; Claessen; Winfield; Stenseth; Vollestad | Proc R Soc Ser B-Bio | 2010 | 10.1098/rspb.2009.1724 |  |
| Mugabo; Perret; Legendre; Le Galliard | J Anim Ecol | 2013 | 10.1111/1365-2656.12109 |  |
| Szostek; Schaub; Becker | J Anim Ecol | 2014 | 10.1111/1365-2656.12206 |  |
| Smith; Cubaynes; MacNulty; Stahler; Quimby; Coulson | Popul Ecol | 2014 | 10.1111/1365-2656.12238 | No MPM |
| O'Farrell; Salguero-Gomez; van Rooij; Mumby | J Anim Ecol | 2015 | 10.1111/1365-2656.12399 | No MPM |
| Crawford; Maerz; Nibbelink; Buhlmann; Norton | J Appl Ecol | 2013 | 10.1111/1365-2664.12194 |  |
| Monadjem; Wolter; Neser; Kane | Anim Conserv | 2013 | 10.1111/acv.12054 |  |
| Keith; Mahony; Hines; Elith; Regan; Baumgartner; Hunter; Heard; Mitchell; Parris; Penman; Scheele; Simpson; Tingley; Tracy; West; Ak√ßakaya | Conserv Biol | 2014 | 10.1111/cobi.12234 |  |
| Ryberg; Hill; Painter; Fitzgerald | Conserv Biol | 2014 | 10.1111/cobi.12429 |  |
| Ciblis-Stewart; Sandercock; McCornack | Entomol Exp Appl | 2015 | 10.1111/eea.12325 |  |
| Kvalnes; Saether; Haanes; Roed; Engen; Solberg | Evolution | 2016 | 10.1111/evo.12952 |  |
| Fordham; Mellin; Russell; Akcakaya; Bradshaw; Aiello-Lammens; Caley; Connell; Mayfield; Shepherd; Brook | Glob Change Biol | 2013 | 10.1111/gcb.12289 |  |
| Riegl; Purkis | Glob Change Biol | 2015 | 10.1111/gcb.13014 |  |
| Altwegg; Jenkins; Abadi | Ibis | 2013 | 10.1111/ibi.12125 |  |
| Sim; Rebecca; Ludwig; Grant; Reid | J Anim Ecol | 2011 | 10.1111/j.1365-2656.2010.01750.x | No MPM |
| Radchuk; Turlure; Schtickzelle | J Anim Ecol | 2012 | 10.1111/j.1365-2656.2012.02029.x |  |
| Johnson; Mills; Wehausen; Stephenson | Ecology | 2010 | 10.1111/j.1365-2664.2010.01846.x | No MPM |
| Gamelon; Gaillard; Servanty; Gimenez; Toigo; Baubet; Klein; Lebreton | J Appl Ecol | 2012 | 10.1111/j.1365-2664.2012.02160.x |  |
| Harris; Fordham; Mooney; Pedler; Araujo; Paton; Stead; Watts; Akcakaya; Brook | J Appl Ecol | 2012 | 10.1111/j.1365-2664.2012.02163.x |  |
| Brodie; Muntifering; Hearn; Loutit; Loutit; Brell; Uri-Khob; Leader-Williams; Preez | Anim Conserv | 2011 | 10.1111/j.1469-1795.2010.00434.x |  |
| Gamelon; Besnard; Gaillard; Servanty; Baubet; Brandt; Gimenez | Evolution | 2011 | 10.1111/j.1558-5646.2011.01366.x |  |
| Di Minin; Griffiths | Ecography | 2011 | 10.1111/j.1600-0587.2010.06263.x |  |
| Hebblewhite; Merrill | Oikos | 2011 | 10.1111/j.1600-0706.2011.19436.x |  |
| DeLong; Melin; Laake; Morris; Orr; Harris | Mar Mammal Sci | 2017 | 10.1111/mms.12427 | No MPM |
| Wang; Ye; Li; Huo; Li; Yu | Restor Ecol | 2016 | 10.1111/rec.12409 |  |
| Boveng; Hoef; Withrow; London | Risk Anal | 2018 | 10.1111/risa.12988 |  |
| Seignobosc; Hemerik; Koelewijn | Int J Ecol | 2011 | 10.1155/2011/870853 |  |
| Sabat; Toledo-Hernandez | J Marin Biol | 2015 | 10.1155/2015/987060 |  |
| Zuniga-Vega; Molina-Zuluaga; Hernandez-Gallegos; Manriquez-Moran; Rodriguez-Romero; Villagran-Santa Cruz; Mendez-de la Cruz | Amphibia-Reptilia | 2012 | 10.1163/15685381-00002843 | Couldn't access: paywall |
| Klok; van Turnhout; Willems; Voslamber; Ebbinge; Schekkerman | Anim Biol | 2010 | 10.1163/157075610X523260 | Couldn't access: paywall |
| Wilson; Martin | BMC Ecol | 2012 | 10.1186/1472-6785-12-9 |  |
| Hostetler; Kneip; Van Vuren; Oli | PLOS ONE | 2012 | 10.1371/jourNAl.pone.0034379 |  |
| Rota; Millspaugh; Rumble; Lehman; Kesler | PLOS ONE | 2014 | 10.1371/jourNAl.pone.0094700 |  |
| Paez; Bock; Espinal-Garcia; Rendon-Valencia; Alzate-Estrada; Cartagena-Otalvaro; Heppell | Copeia | 2015 | 10.1643/CE-14-191 |  |
| Corti; Wittmer; Festa-Bianchet | J Mammal | 2010 | 10.1644/09-MAMM-A-047.1 |  |
| Zamora-Abrego; Chang; Zuniga-Vega; Nieto-Montes de Oca; Johnson | Herpetologica | 2010 | 10.1655/09-005.1 |  |
| Perez-Mendoza | Herpetologica | 2013 | 10.1655/HERPETOLOGICA-D-12-00038R2 |  |
| Rigby; Haukos | Southeast Nat | 2014 | 10.1656/058.013.s505 |  |
| Bulluck; Buehler; Vallender; Robertson | Wilson J Ornithol | 2013 | 10.1676/12-154.1 |  |
| Sakaris; Irwin | Ecol Appl | 2010 | 10.1890/08-0305.1 |  |
| McMurray; Henkel; Pawlik | Ecology | 2010 | 10.1890/08-2060.1 |  |
| Hunter; Caswell; Runge; Regehr; Amstrup; Stirling | Ecology | 2010 | 10.1890/09-1641 |  |
| Carson; Cook; Lopez-Duarte; Levin | Ecology | 2011 | 10.1890/11-0488.1 |  |
| Bjorkvoll; Lee; Grotan; Saether; Stien; Engen; Albon; Loe; Hansen | Ecology | 2016 | 10.1890/15-0317.1 |  |
| Hernandez-Pacheco; Hernandez-Delgado; Sabat | Ecosphere | 2011 | 10.1890/ES10-00065.1 |  |
| Grear; Meyer; Cooley; Kuhn; Piper; Mitro; Vogel; Taylor; Kenow; Craig; Nacci | J Wildlife Manage | 2010 | 10.2193/2008-093 |  |
| Mogollones; Rodroguez; Hernandez; Barreto | Chelonian Conserv Bi | 2010 | 10.2744/CCB-0778.1 |  |
| Suzuki; Kobayashi; Nakamura; Takasu | Wildlife Biol | 2013 | 10.2981/13-021 |  |
| Whitehead; Gero | Endanger Species Res | 2015 | 10.3354/esr00657 |  |
| Linares; Doak | Mar Ecol Prog Ser | 2010 | 10.3354/meps08437 |  |
| Kelly; Metaxas | Mar Ecol Prog Ser | 2010 | 10.3354/meps08442 |  |
| Edmunds | Mar Ecol Prog Ser | 2010 | 10.3354/meps08595 | No MPM |
| Button; Rogers-Bennett | Mar Ecol Prog Ser | 2011 | 10.3354/meps09094 | No MPM |
| Jimenez-Melero; Ramirez; Guerrero | Freshwater Biol | 2013 | 10.3354/meps10377 |  |
| New; Clark; Costa; Fleishman; Hindell; Klanjscek; Lusseau; Kraus; McMahon; Robinson; Schick; Schwarz; Simmons; Thomas; Tyack; Harwood | Mar Ecol Prog Ser | 2014 | 10.3354/meps10547 |  |
| Ali; Kauffman; Amin; Kibara; King; Mallon; Musyoki; Goheen | Ecol Appl | 2018 | 10.5061/dryad.480tf |  |
| Bijlsma; Vermeulen; Hemerik; Klok | Ardea | 2012 | 10.5253/078.100.0208 |  |
| Beissinger | PeerJ | 2014 | 10.7717/peerj.549 |  |
| Watsa | NA | 2013 | 10.7936/K7DB7ZTD | Not peer reviewed |
| Velez-Espino; Ford; Araujo; Ellis; Parken; Balcomb | Can Tech Report Fish & Aq Sci | 2014 | 978-1-100-23563-9 | Not peer reviewed |
| Defler | NA | 2014 | 978-1-4939-0697-0 |  |
